# Gentrius: identifying equally scoring trees in phylogenomics with incomplete data

**DOI:** 10.1101/2023.01.19.524678

**Authors:** Olga Chernomor, Christiane Elgert, Arndt von Haeseler

## Abstract

Phylogenetic trees are routinely built from huge and yet incomplete multi-locus datasets often leading to phylogenetic terraces – topologically distinct equally scoring trees, which induce the same set of per locus subtrees. As typical tree inference software outputs only a single tree, identifying all trees with identical score challenges phylogenomics. Generating all trees from a terrace requires constructing a so-called stand for the corresponding set of induced locus subtrees. Here, we introduce Gentrius – an efficient algorithm that tackles this problem for unrooted trees. Despite stand generation being computationally intractable, we showed on simulated and biological datasets that Gentrius generates stands with millions of trees in feasible time. Depending on the distribution of missing data across species and loci and the inferred phylogeny, the number of equally optimal terrace trees varies tremendously. The strict consensus tree computed from them displays all the branches unaffected by the pattern of missing data. Thus, Gentrius provides an important systematic assessment of phylogenetic trees inferred from incomplete data. Furthermore, Gentrius can aid theoretical research by fostering understanding of tree space structure imposed by missing data.

**One-Sentence Summary:** Gentrius - the algorithm to generate a complete stand, i.e. all binary unrooted trees compatible with the same set of subtrees.

## Main Text

In molecular phylogenomics one infers a tree for a group of species using genetic information from many different loci. Contemporary datasets often comprise hundreds of species and hundreds of loci, combining organisms from distant taxonomic groups, and well-studied species, whose genomes were sequenced completely, together with non-model organisms, for which only a handful of loci are available. As a result, such diverse datasets often exhibit missing data, i.e. for some species, sequences for some loci are not available, either as a consequence of species-specific gene losses/acquisitions or incompletely sequenced genomes. The availability of genetic sequences is summarised in a species per locus presence-absence matrix with 1’s or 0’s indicating presence or absence of a sequence, respectively (e.g., Fig. 1A). Typical percentage of zeros (i.e. missing data) in such matrices ranges from 30% to 80% (e.g. Table 1). Missing data affects phylogenetic methods in various ways (e.g. ^1–7^).

**Fig. 1.**
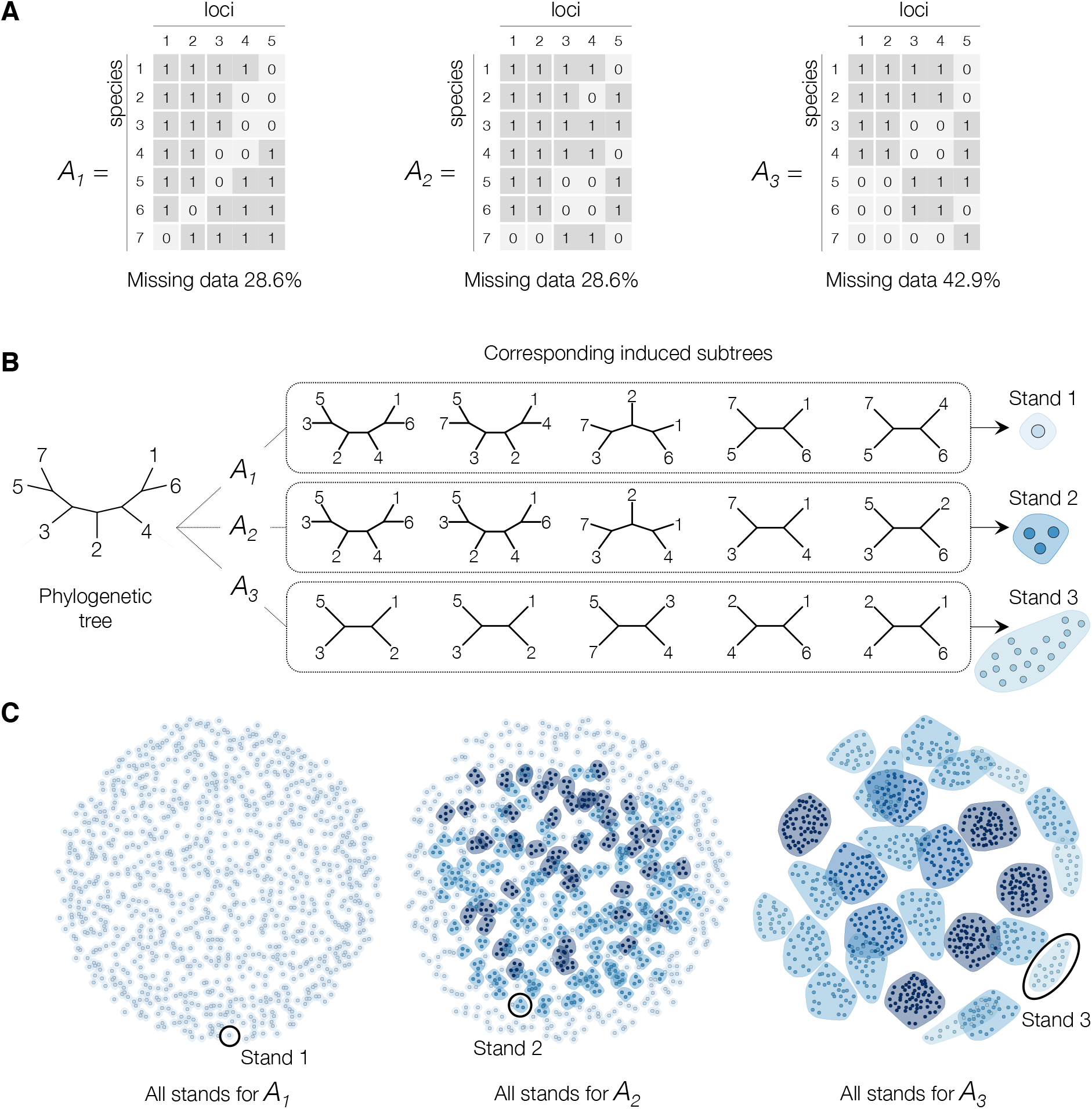
Influence of missing data on phylogenomic inference. (**A**) Examples of species per locus presence-absence matrices. Here, “1” stands for presence and “0” for absence of sequence for corresponding species and locus. (**B**) A random tree with its induced subtrees and corresponding stands for each matrix. Each dot in a stand is a different tree, i.e. stands 1, 2, and 3 consist of 1, 3 and 17 trees, respectively. (**C**) For each considered matrix all 945 trees for seven species were grouped based on their induced subtrees into stands.

**Table 1.** Summary for biological datasets. Column “Max locus coverage” for a given dataset indicates information about the largest number of species in one locus (i.e. the largest column sum in a presence-absence matrix). Column “Sp. min coverage” contains the number of species, represented by a single locus, thus have minimal coverage (i.e. row sum equals to one). The lower bound on stand size for partially generated stands is 100M trees.

| ID | Species group | Sp. | Loci | Missing data | Max locus coverage | Sp. min coverage | Stand size | Total CPU time |
| --- | --- | --- | --- | --- | --- | --- | --- | --- |
| D1 | Snails<sup>34</sup> | 62 | 1027 | 37% | 58 (94%) | 0 | 1 | 2s |
| D2 | Land plants<sup>35</sup> | 103 | 852 | 34% | 100 (97%) | 0 | 1 | 2s |
| D3 | Drosophilinae<sup>36</sup> | 180 | 15 | 60% | 144 (80%) | 6 | 127,575 | 5s |
| D4 | Carnivora<sup>37</sup> | 237 | 74 | 75% | 202 (85%) | 22 | 29,133 | 6s |
| D5 | Frogs<sup>38</sup> | 267 | 20 | 59% | 266 (99.6%) | 3 | 1 | 0.04s |
| D6 | Primates-1<sup>39</sup> | 279 | 27 | 74% | 202 (72%) | 34 | 9,963 | 9s |
| D7 | Grasses<sup>40</sup> | 298 | 3 | 34% | 221 (74%) | 113 | >100 M | 26m:24s |
| D8 | Primates-2<sup>41</sup> | 372 | 79 | 63% | 276 (74%) | 57 | 39,355,875 | 1h:20m:3s |
| D9 | Salamander<sup>42</sup> | 381 | 13 | 60% | 314 (82%) | 77 | >100 M | 1h:22m:40s |
| D10 | Monocots<sup>11</sup> | 404 | 11 | 63% | 290 (72%) | 45 | 14,529,375 | 10m:53s |
| D11 | Sedges<sup>43</sup> | 435 | 18 | 80% | 329 (76%) | 93 | >100 M | 7h:39m:22s |
| D12 | Snakes<sup>44</sup> | 767 | 5 | 48% | 716 (93%) | 59 | 315 | 0.064s |

One of the crucial impacts of missing data on tree inference are phylogenetic terraces – topologically distinct trees, which due to combinatorial structure and the considered scoring function, have equal scores^1^. Terraces arise for parsimony and likelihood^1,2^, as well quartet consistency^8^. Thus, they affect two popular tree inference approaches: supermatrix and supertree. Terraces influence the efficiency of tree searches^9–11^ and introduce biases to bootstrap^2^. Finally, if a best-found tree is on a large terrace^1^, ignoring this fact may lead to overconfident conclusions about evolutionary relationships^1,8,12^. Thus, the knowledge of all trees on a terrace will improve the reliability of a phylogenetic analysis.

Please note, that even for complete datasets multiple equally scoring trees can occur (e.g. well-known for parsimony and supertree methods^13–15^). In such cases the researcher has little influence on the scoring landscape. In contrast, phylogenetic terraces are caused by missing data, which can be tampered with to minimise the effect on phylogenetic inference.

How do we determine the size of a terrace and trees on it (of note, can be exponentially many^16^)? The trees from a terrace have a well-defined combinatorial structure: they all share (i.e. compatible with) the same set of induced per locus subtrees. Here, each induced subtree is obtained from a tree (any terrace tree) by removing species with zeros for the corresponding locus in the presence-absence matrix (Fig. 1B). Thus, generating terrace trees is equivalent to generating a so-called stand^2^ - a collection of all trees, which are compatible with the (induced) subtrees.

Unfortunately, for unrooted binary subtrees, constructing a stand or determining that it is empty (i.e. the subtrees are incompatible) is computationally intractable^17^. Current methods^1,18^, adaptations from the algorithm for rooted trees^19^, are limited to a special case. Other theoretical work focused on the existence of multiple trees^20^ on the stand with neither generating nor enumerating them all and thus not allowing any post-analysis.

To close this gap and to provide a practically useful solution, we developed Gentrius - a deterministic branch-and-bound type algorithm to generate complete stands from binary unrooted subtrees. Applied to phylogenetic terraces, for a tree inferred with any phylogenomic method (e.g. IQ-TREE 2^21^, RAxML^22^, ASTRAL^23^ etc.) and a species per locus presence-absence matrix, Gentrius generates all trees from the corresponding stand (thus, terraces for parsimony, likelihood, quartet consistency score). Hence, as terraces are equally scoring trees due to missing data, Gentrius systematically assesses the influence of missing data on their number and topologies and enhances the confidence of evolutionary conclusions one may draw from the data. When all trees from a stand/terrace are generated, one can subsequently study their topological differences employing routine phylogenetic approaches. Moreover, we also provide a handy summary and visualisation script to ease the post-analysis for non-technical experts. To foster a widespread application, Gentrius was implemented in IQ-TREE 2^21^. In the following we elucidate the impact of missing data on stand size, briefly describe the Gentrius algorithm, test its feasibility on simulated data, and show application to empirical data identifying phylogenetic terraces equipped with likelihood scoring function. We conclude by discussing approaches to assure the robustness of phylogenomic analysis in the presence of missing sequences.

## Results

### On missing data and stands

To illustrate the influence of missing data on phylogenomic inference, we first discuss three fictive examples (Fig. 1A) with seven species, five loci and presence-absence matrices *A*_1_, *A*_2_, *A*_3_ with 29%, 29%, and 43% of missing data, respectively. For each matrix and each of the possible 945 binary unrooted seven species trees we computed induced subtrees (Fig. 1B) and partitioned the 945 trees with the same set of induced subtrees into stands (Fig. 1C). For matrix *A*_1_ each stand contained one tree. Matrix *A*_2_, with 29% of missing data like *A*_1_, leads to 360 stands of size one and 165 stands comprising two or more trees. This indicates that stand sizes depend not only on the percentage, but also on the spread of zeros in the presence-absence matrix. Finally, for matrix *A*_2_, with the largest percentage of missing data (43%), stand sizes varied from 17 to 59 trees. Thus, predicting the stand size from the presence-absence matrix alone is challenging and requires appropriate computational approaches, which motivated development of Gentrius.

### Gentrius in a nutshell

For simplicity, we explain the modus operandi of Gentrius with an illustrative example (Fig. 2) and for details see Supplementary Note 1. To generate a stand Gentrius uses the information provided by the (induced) subtrees (Fig. 2A, B). It selects one of the subtrees as initial subtree (black subtree in Fig. 2B) and then inserts on it the missing species (3, 4, and 7) sequentially, starting with species 3. The initial subtree has nine branches and, thus, nine possibilities to insert species 3. The remaining input subtrees (red and blue) constrain the placement for insertion. The red subtree and the initial subtree have species 1, 2, and 9 in common (Fig. 2C left). Further, according to the red subtree species 1 and 9 are more closely related in contrast to 2 and 3. Thus, to preserve this relationship, it is not allowed to insert species 3 on the path connecting 1 and 9 in the initial subtree, allowing only seven branches for insertion (red dots in Fig. 2C left). Similarly, to agree with the blue subtree, species 3 can be inserted only on five branches (blue dots in Fig. 2C middle). Finally, their intersection, the three branches with red/blue dots are allowed by both constraint subtrees (Fig. 2C right). They constitute admissible branches for insertion of species 3. Inserting species 3 on them generates three new intermediate subtrees (Fig. 2D, second row). This procedure is continued with the next missing species and each intermediate subtree (Fig. 2D, third row) until all missing species are inserted generating all seven trees from the corresponding stand.

**Fig. 2.**
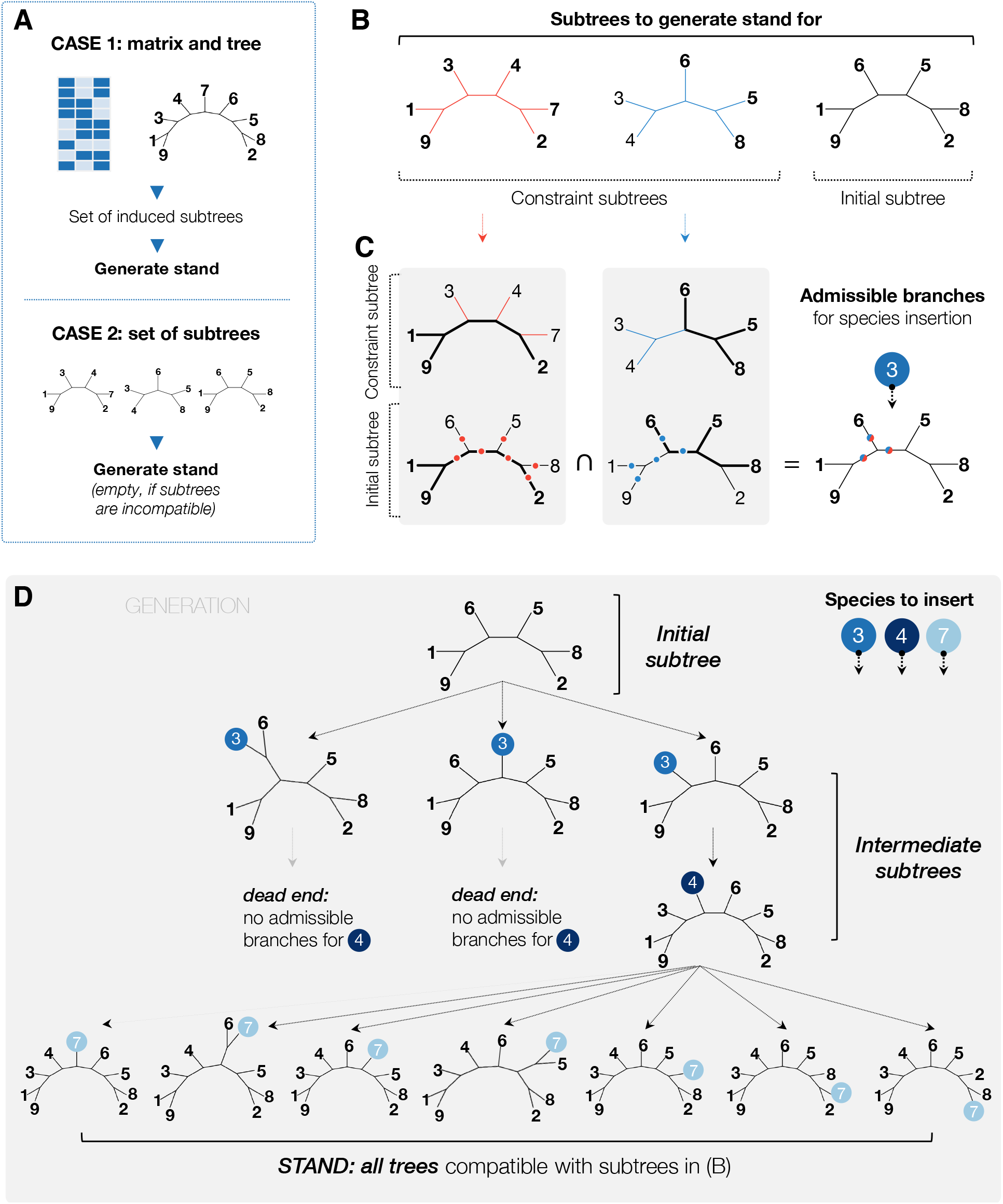
The overview of Gentrius exemplified on an small dataset. (**A**) Gentrius generates stands from a set of binary unrooted subtrees. They can be obtained either as induced subtrees of a tree and a presence-absence matrix (CASE 1, anticipated practical application in a typical phylogenomic workflow), or as a set of subtrees inferred separately (CASE 2). (**B**) The set of subtrees to generate a stand for. The black subtree is selected as initial subtree. The remaining subtrees (red and blue) serve as constraints. (**C**) Identification of admissible branches to insert species 3. First, we detect branches, which are allowed by each constraint subtree separately (left, middle) and then compute their intersection (right). Bold black branches connect species in common for corresponding pairs of initial and constraint subtree. Red, blue and red-blue dots mark branches on initial subtree allowed by red, blue and both constraint subtrees, respectively. (**D**) Generation of a stand. Species 3, 4 and 7 are inserted sequentially. Each insertion generates an intermediate subtree. After species 3 is inserted, Gentrius identifies admissible branches for species 4. If there are no admissible branches, a dead end is reached, i.e. the intermediate subtree cannot be extended without constraint violation, and Gentrius continues with the next intermediate subtree. After species 4 is inserted, we identify the admissible branches for species 7. By iterating over all admissible branches and inserting all missing species Gentrius generates a complete stand, i.e. all trees compatible with an input set of subtrees in (B).

To detect admissible branches we developed edge maps (Supplementary Note 2, Extended Data Fig. 1) and data structure (Supplementary Note 3) that allow considering all subtrees simultaneously. This is necessary for detecting all admissible branches, which is the core of Gentrius and assures generating the stand completely. The efficient identification of admissible branches and their updating during species insertion (and deletion), as well as the choice of the initial subtree and the insertion order of missing species are all crucial considerations for the performance of Gentrius.

However, the number of trees on the stand can be exponential^16^ and in such cases generating a complete stand in feasible time is not possible. Therefore, Gentrius employs a user-defined threshold, MaxStandTrees (for details see Supplementary Note 3), on the maximal number of trees generated from a stand. If MaxStandTrees is reached, Gentrius outputs a partial stand. Moreover, the maximum number of intermediate subtrees is bounded by the threshold MaxIntermediate, since also the number of intermediate subtrees can be exponential^17^. In the worst case MaxIntermediate might be reached even before any tree from a stand is generated. To tackle such computationally complex cases, we provide an alternative setting for the initial subtree (Supplementary Note 1.2), which comes with a limitation that Gentrius generates a stand only partially. The above cases define different difficulty levels for Gentrius: a stand is generated completely; partially with tree number equal to MaxStandTrees; partially, triggering MaxIntermediate with either stand size > 0 or empty stand (i.e. complex dataset). We consider the application of Gentrius to a given data as successful, if either Gentrius generates a stand completely, or partially, thus, providing a lower bound on the stand size.

### Performance on simulated data

To evaluate the feasibility of Gentrius we performed an extensive simulation study with varying species number (20-700), locus number (5-100), different percentages (30%, 50%, 70%) and coverage patterns of missing data. To simulate different coverage patterns we developed a custom matrix simulator (Supplementary Note 4) and controlled the distribution of zeros across matrix via various parameters (Methods). We simulated in total 6,120 presence-absence matrices and for each matrix sampled five random trees^24^, thereby generating 30,600 pairs of presence-absence matrix and tree. For each instance we ran Gentrius with MaxStandTrees and MaxIntermediate set to 100 million (M) trees each.

Figure 3 and Extended Data Figure 2 summarise key aspects of the simulation. Importantly, our simulation-instances covered all complexity levels of the input (listed in the previous section). Despite varying complexity all runs finished in reasonable time (from milliseconds up to ∼31 hours, Fig. 3A, Extended Data Fig. 2A, Supplementary Table 2). In contrast to classical tree *inference* methods (i.e. supertree and supermatrix), which depend a lot on the species and locus numbers, Gentrius runtime predominantly depends on the stand size and, thus, on the complexity level of the input data (Fig. 3A, Extended Data Fig. 2A). If a run stopped reaching MaxIntermediate without generating any tree from the stand (i.e. highest complexity level), we re-run Gentrius employing the alternative initial subtree approach (Supplementary Note 1). In all such cases, the alternative approach generated a partial stand with 100M trees. Thus, Gentrius successfully tackled all instances providing either a complete or a partial stand.

**Fig. 3.**
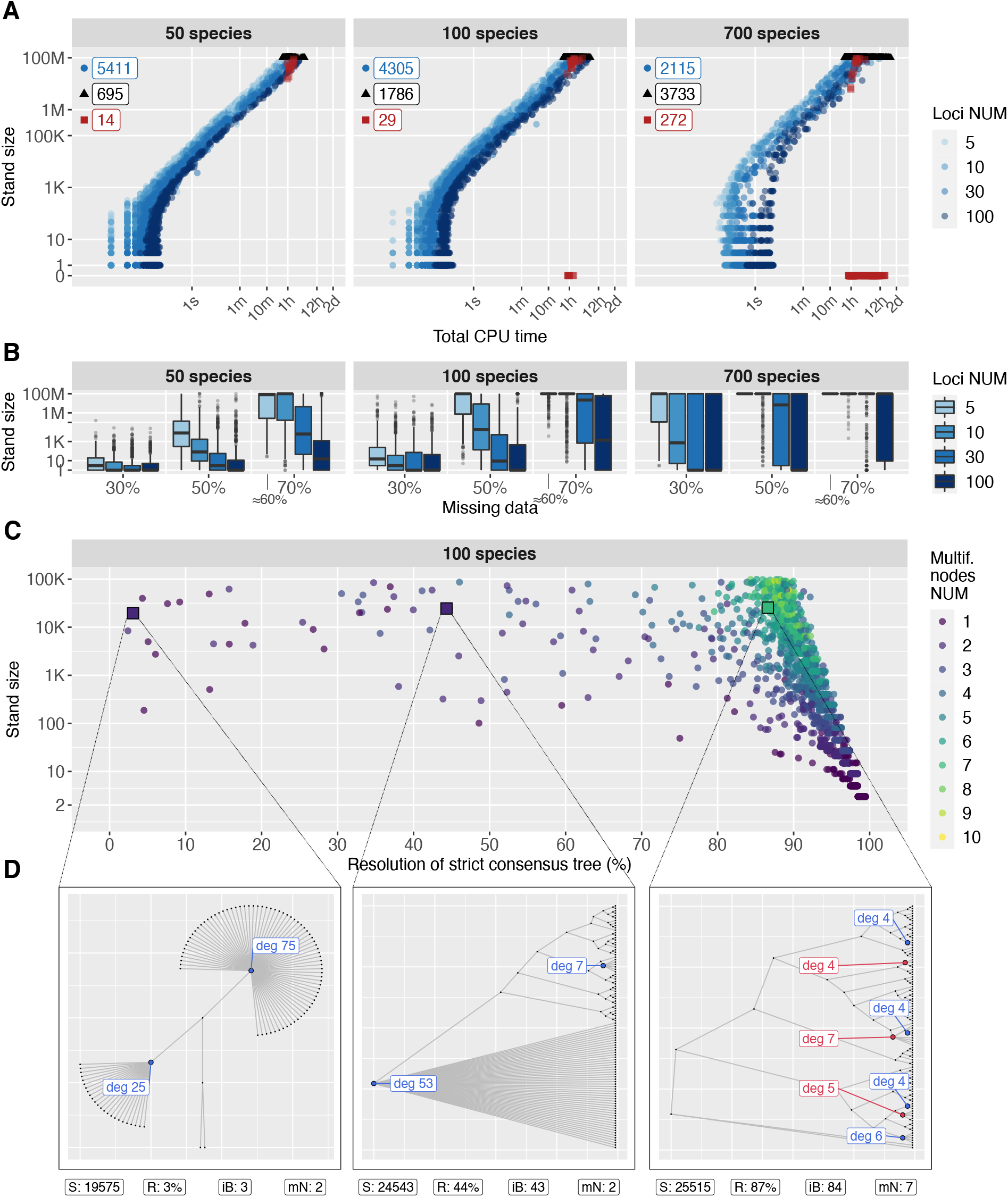
Results of simulation study. (**A**) Stand size vs. total CPU runtime. Shapes correspond to different computational cases: circles denote complete stands, triangles and squares denote incomplete stands, i.e. triggering MaxStandTrees or MaxIntermediate, respectively. The numbers (top left) specify how often each case occurred. The squares at zero represent complex cases, where MaxIntermediate was triggered, but no tree from a stand was generated yet. For all datasets with red squares, we subsequently ran Gentrius with an alternative initial subtree (Methods), confirming a lower bound of 100M trees on the stand size. (**B**) Influence of the percentage of missing data on stand sizes. Note, that for 5 loci with the constrains imposed (Methods), the resulting presence-absence matrices had around 60% of missing data instead of 70% as required (Supplementary Fig. 4). (**C**) Resolution of strict consensus trees for 100 species. Each point corresponds to one stand. All stands have up to 100K trees. The resolution refers to the percentage of internal branches on a strict consensus tree compared to the *n* − 3 internal branches on a fully resolved tree with *n* species. (**D**) Selected examples of strict consensus trees with different resolution. Multifurcating nodes are marked on the tree by coloured circles and degree of the node. Blue nodes are incident to exactly one internal branch and red nodes are incident to at least two internal branches. Large node degrees represent highly unresolved parts of the tree due to missing data. S – stand size, R – tree resolution, iB – number of internal branches, mN – number of multifurcating nodes.

Finally, we compared the runtimes of Gentrius with Terraphast^18^, that generates stands only if the data contains at least one reference species (i.e. a species with no missing data). This condition together with the fact that we had to limit the stand size to less than 100M trees to compare the computing times, left us with 10,293 simulated instances. Overall, the runtimes of both software is very similar (Extended Data Fig. 3). Notably, Gentrius tends to be faster, when the species number is larger than locus number. The biggest advantage of Gentrius, however, is that it does not impose any requirements on the input data.

### Topological differences

In a phylogenomic analysis, all trees belonging to the phylogenetic terrace of the highest scoring tree are plausible hypotheses about the relationship of organisms (i.e. equally optimal trees). Thus, generating terraces is important to understand the influence of missing data on phylogenetic relationships and to elucidate reliable branching patterns in the multiple best found trees.

To investigate the similarity of trees from the same terrace, we analysed trees from 8,440 stands with stand sizes between 2 and 100K and with 20-100 species. For each stand we computed the strict consensus tree to identify common branching patterns. Here, the strict consensus tree displays branches occurring in all trees from a stand. A fully resolved binary tree with *n* species has *n* − 3 internal branches, while the occurrence of multifurcating nodes (i.e. nodes with more than three outgoing branches) reduces the number of internal branches and, thus, tree resolution. To measure the loss of resolution (here, caused by missing data) for each strict consensus tree we computed the percentage of its internal branches compared to *n* − 3 internal branches on a fully resolved tree.

Figure 3C shows the relation between resolutions of the strict consensus trees and stand size for 2,067 stands for datasets with 100 species (see also Extended Data Fig. 4). We observed the full range of resolutions (between 3%-99% and up to 10 multifurcating nodes), where 93% of the consensus trees showed a resolution ≥ 85%. Importantly, we observed that the resolution cannot be predicted from the stand size. For instance, the strict consensus trees for stands with similar sizes (from ∼20K to ∼26K trees) show a resolution of 3%, 44% and 87% (Fig. 3D, see also Extended Data Fig. 4).

Therefore, for non-trivial phylogenetic terraces (i.e. stand size > 1), it is crucial to know all its trees to precisely compute the amount and location in a tree of phylogenetic uncertainty due to missing data. Note, that this uncertainty should not be mixed with bootstrap support^25^, which is, in fact, also affected by missing data^2^.

Note that, while a strict consensus tree is a good measure for the congruence/compatibility of trees from a terrace/stand, it should only be used if a stand is generated completely. A partial stand even with millions of trees would not be representative for potentially an exponentially large stand. Even more so, since consecutive trees generated by Gentrius are more similar to each other. Thus, if Gentrius outputs a partial stand, the researcher must increase the stopping thresholds and generate a complete stand, before constructing strict consensus tree.

### Practical advice to reduce stand size

Knowing which presence-absence matrices lead to huge stands would help designing appropriate datasets before performing computationally intensive phylogeny inference and then evaluating the impact of missing data and terraces. However, predicting stand size even for a specific tree is generally not feasible. One trivial case is when at least one locus has no missing data, then stands for all trees have size one (i.e. only trivial terraces). However, complete loci are rare in empirical datasets with large number of taxonomically diverse species.

In Figure 3B we showed that stand size depends on the % of missing data, species and locus numbers. However, simply increasing the number of loci does not necessarily reduce the stand size (e.g. results for 700 species in Fig. 3B). As exemplified in Fig. 1, stands sizes also depends on the spread of missing data across the presence-absence matrix.

To get a better understanding, we simulated presence-absence matrices with different coverage pattern, but fixed the amount of missing data across either species or loci to uniform.

Figures 4A-C show the effect of presence-absence matrices if their locus coverage pattern changes, while the amount pf missing data across species is uniform. Fig. 4B shows the changing locus coverage pattern from a “uniform” distribution of missing data (horizontal line) to different bi-modal distributions (step-type functions). The amount of missing data across species is fixed (i.e. uniform coverage). Not surprisingly having a few densely sampled loci reduces the stand size.

**Fig. 4.**
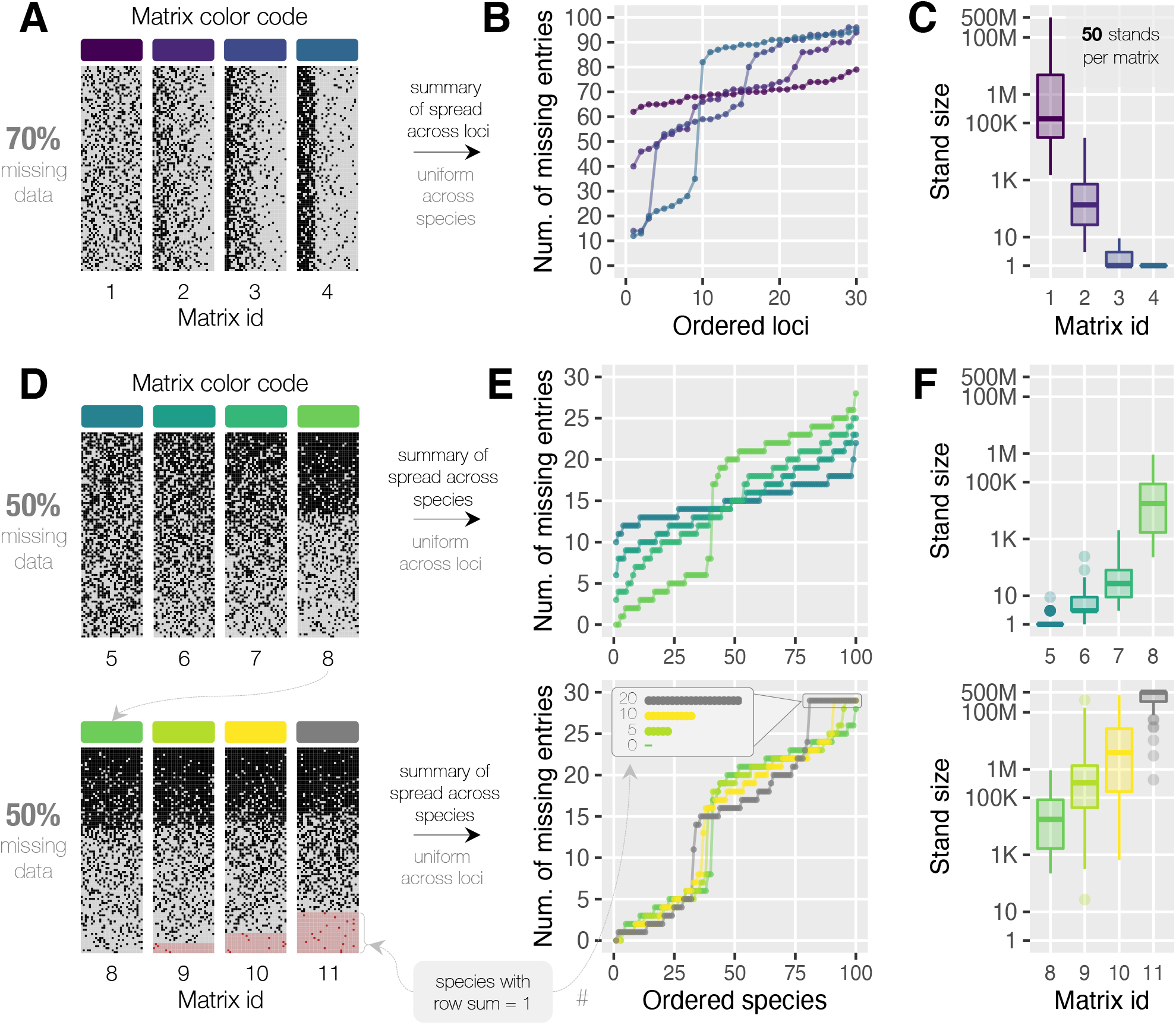
Dependence of stand size on missing data across species and loci. The simulated presence-absence matrices have 100 species and 30 loci. Black and grey dots indicate presence or absence (“1” or “0”) of sequence for corresponding species and locus. For stand size computation the 50 trees were fixed across all matrices. (**A**) Matrices with 70% of missing data. Each species has exactly 21 missing entries, while spread of missing data across loci varies among matrices. (**B**) For each matrix from (A) the loci are ordered by increasing missing data. (**C**) Stand sizes for matrices from (A). (**D**) Matrices with 50% of missing data. Each locus has exactly 50 missing entries, while spread of missing data across species varies among matrices. Species represented by a single locus are marked by pink row with a red dot indicating presence of sequence. Such minimally covered species are only present in matrices 9, 10, and 11 which have 5, 10 and 20 minimally covered species respectively. (**E**) For each matrix from (D) the species are ordered by increasing missing data. (**F**) Stand sizes for matrices from (D).

Figures 4D-F show the complementary analysis. Now, the species coverage pattern changes and the amount of per locus missing data is fixed. Note, that some species here are only represented by a single locus. Now, the number of trees on a stand increases as the deviation of the missing data from the uniform distribution increases. Thus, species with sparse data increase the stand size. For other sizes of matrices see also Extended Data Figs. 5, 6.

Thus, stand/terrace sizes are likely to be small, when the missing data per species are uniformly distributed and if at least some loci are very well sampled. For real data, it can be helpful to exclude sparsely sampled species and if possible, sequence/collect additional loci to increase the coverage for some loci. For the latter case Gentrius helps to systematically evaluate, which changes in the presence-absence matrix will lead to a size reduction of the stand/terrace without too much extra work or without excluding too many data.

### Application to biological data and phylogenetic terraces with likelihood

The direct practical application of Gentrius is identifying phylogenetic terraces – equally scoring trees due to missing data. Here we will analyse phylogenetic terraces for likelihood inference with edge unlinked partition model^1,2^. Namely, we will generate a terrace for maximum likelihood (ML) tree, thus, identifying all equally optimal ML trees for each dataset. The main questions are how many such optimal trees per dataset there are (i.e. terrace size) and how similar their topologies. This will give us a quantitative answer, if missing data hampers phylogenetic analysis via multiple equally scoring trees and, thus, if the quality of the input dataset is sufficient to make reliable phylogenetic statements.

To this end we analysed 12 published alignments (Table 1, Supplementary Fig. 5) from various taxonomic groups. For each alignment, its corresponding presence-absence matrix and inferred ML tree (Methods) we generated the stand/terrace with Gentrius.

In accordance with our simulations, the resulting stand sizes were highly variable (Table 1), ranging from one (for Snails, Land plants and Frogs) to more than 100M trees (for Grasses, Salamander and Sedges). We would like to stress that the results anew support the conclusion from simulations (Fig. 3A), that the computing times (Table 1) are by and large determined by the stand size, rather than locus number.

In the simulations we have observed that stand sizes for the same presence-absence matrix vary a lot for different trees (Fig. 4C,F, Extended Data Figures 5D, 6D). To confirm this on empirical presence-absence matrices, we additionally sampled 100 random trees for each alignment and generated their terraces. For Snails and Land plants all terraces have size one (Supplementary Table 4). This is expected, since both datasets contain many loci with few missing sequences (Supplementary Fig. 5). For Frogs dataset terrace sizes varied between one to seven trees. Finally, for nine datasets the majority of terraces have more than 100M trees. This observation corroborates our simulations, that it is difficult to predict the stand/terrace size from a presence-absence matrix alone.

For six datasets with complete terraces (for ML trees) with sizes from 315 to ∼40M trees, we computed their strict consensus trees. Despite generally high resolution of the strict consensus trees (from 99% for Snakes to 78% for Carnivora, Fig. 5, and Supplementary Fig. 6-11), for a careful interpretation of the results, the characteristics of each multifurcation, as indicators of uncertain branching patterns due to missing data are important.

**Fig. 5.**
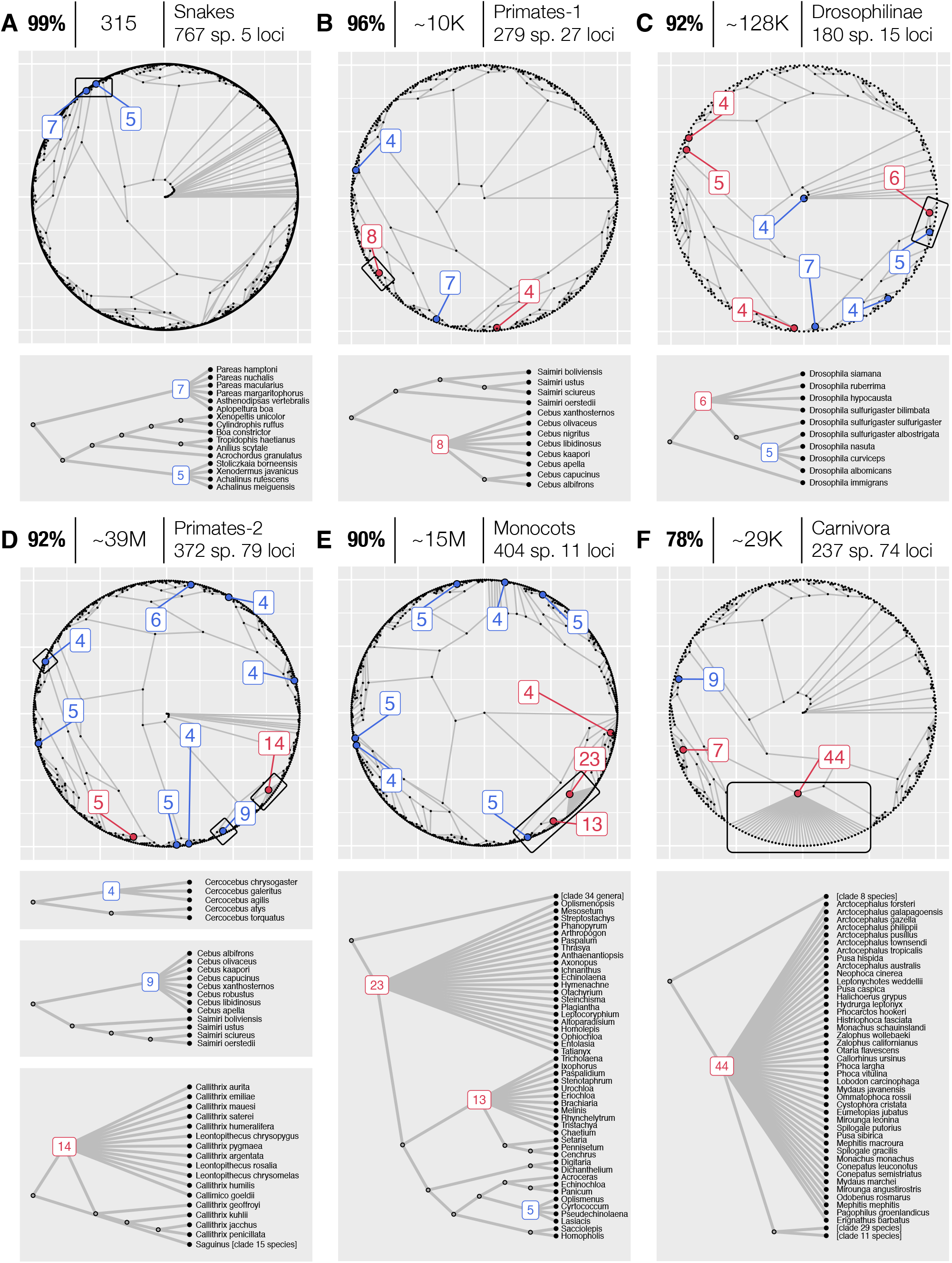
Strict consensus trees for (complete) stands generated for ML trees for six biological datasets. The subplots are ordered by the decreasing resolution (in %) of strict consensus trees (left), followed by stand size (middle), species group, numbers of species and loci (right). The numbers in blue and red indicate the degree of multifurcation, where blue nodes are incident to exactly one internal branch and red nodes are incident to at least two internal branches. Black boxes on the circular trees denote the subtrees displayed below each subplot. These subtrees were chosen to illustrate different cases of uncertainties (different types of nodes (red/blue), node degree, species involved). (**A**) Strict consensus tree for Snakes with 48% of missing data. (**B**) Primates-1, 74%. (**C**) Drosophilinae, 60%. (**D**) Primates-2, 63%. (**E**) Monocots, 63%. (**F**) Carnivora, 75%.

For instance, if a dataset contains species from diverse taxonomic groups and each multifurcating node only affects species from the same genus (e.g. see subtrees in Figs. 5B, D), then one can still make reliable statements about the evolution of genera, even if the number of multifurcations is high. However, if the evolutionary question concerns species from the same genus/family, like in Drosophilinae (Fig. 5C), then multifurcations prohibit a detailed analysis of the phylogenetic relationship of the species. The biggest issue is, of course, when multifurcating nodes involve species from different taxonomic groups and no conclusions can be drawn about their evolution as a result of missing data and terraces (e.g. Fig. 5A,D,E,F).

In general, large number of multifurcating nodes and high node degree are indicators of analysis strongly affected by missing data. The above findings anew demonstrate that for a robust phylogenomic analysis taking into account missing data systematically is very important. Here, we advocate including Gentrius into a phylogenomic workflow to increase the confidence of findings.

## Discussion

The main result of the paper is the development of Gentrius, a deterministic branch-and-bound like algorithm to generate stands for unrooted binary subtrees. The evaluation of Gentrius feasibility on simulated and biological datasets evidenced that Gentrius can deal with various datasets, generating millions of trees within reasonable time. To the best of our knowledge, Gentrius is the first algorithm to generate complete stands for unrooted trees without any constraints on the structure and type of input data.

Importantly, Gentrius has direct practical application generating phylogenetic terraces - equally scoring trees due to missing data. As the number of species with diverse genome composition (e.g. species-specific gene losses and acquisitions) grows, collecting complete datasets with thousands of species is challenging. Therefore, missing data is an intrinsic property of datasets with highly diverse species. Thus, generating terraces and investigating impact of missing data should be routine in phylogenetic workflow to assure reliable phylogenetic inference.

Here, we have demonstrated, that sizes of stands (and, thus, terraces) are strongly affected by missing data. Moreover, trees from the same stand can be topologically very diverse. Therefore, if the stand size is larger than one, it is important to investigate topological differences of stand trees, for instance, by constructing a strict consensus tree and subsequently investigating its unresolved parts. If the evolutionary relationships of interest are not affected by multifurcations, then the strict consensus tree can be used to substantiate evolutionary hypotheses. Otherwise, the results have to be considered with care. Ideally, the input data should be amended and reanalysed.

Huge non-trivial terraces are a sign that the input data is not sufficient for reliable phylogenetic conclusions. However, a terrace size of one does not exclude the existence of other equally or almost equally good trees^26,27^. For instance, note that for supertree methods and parsimony, trees from multiple terraces can have identical score. Thus, generating one stand/terrace for these methods gives only a lower bound on the total number of equally scoring trees.

In likelihood methods all stand trees have identical score (i.e. stand is terrace) under edge unlinked partition model^2^. To avoid terraces, one may employ different partition schemes or more restrictive partition models^2^. Then trees from one stand do not have identical score. However, yet very similar likelihoods^28^. Thus, to generate trees with similar quality for alternative partition models or schemes, instead of expensive multiple runs of likelihood inference, one can use stands, and subsequently analyse stand trees by computing their scores and using tree tests (e.g. ^29^). If missing data are unavoidable, the best strategy is to combine a large number of nearly complete loci and avoiding species with a lot of missing data.

Importantly, phylogenetic terraces and missing data affect not only tree inference, but also bootstrap^25^ – a classical method for assessing clade confidence. Namely, a tree for each bootstrap replicate (i.e. resampled alignment) might lie on a large terrace of its own^2^, which may cause misleading bootstrap scores^2^. Even summarising bootstrap trees becomes non-trivial^2^. Importantly, bootstrap cannot explicitly say anything about the effect of missing data on equally optimal trees. In contrast, by generating all terrace trees for the best found tree Gentrius provides an unbiased deterministic summary of the effect of missing data on the number and topology of equally optimal trees.

Apart from direct practical application to phylogenetic terraces, Gentrius fosters our theoretical understanding of tree spaces (set of all trees for a given species number, e.g. Fig. 1C). For instance, it allows investigating scoring landscapes imposed by missing data as well as studying how trees from the same stand and also between different stands are connected via common topological rearrangements (e.g. Nearest Neighbour Interchange, see also ^30^). This knowledge has a great potential for development of better tree search strategies in the presence of missing data ^9,10^.

## Methods

**Extended description of Gentrius algorithm** (Supplementary Note 1).

**Identifying admissible branches** (Supplementary Note 2).

**Implementation, complexity and validation** (Supplementary Note 3).

**Custom matrix simulator** (Supplementary Note 4).

**Comparison with Terraphast** (Supplementary Note 5).

### Simulated presence-absence matrices and trees

#### SIM1

To test feasibility of Gentrius we simulated 6,120 binary matrices using a custom matrix simulator (Supplementary Note 4). For each triplet of species number (rows), locus number (columns) and the percentage of missing data (zeros in a matrix), we varied spread of zeros across rows and columns. Starting with a matrix of one’s, the zeros were distributed using row and column sampling probabilities, subject to constraints on row/column sums. The sampling probabilities were chosen to include various combinations of rows and columns with high, medium and low amounts of missing data. Basic constraints on row/column sums included: (i) column sum ≥ 4 (i.e. locus covers at least four species); (ii) each column has missing data; (iii) row sum ≥ 1 (i.e. each species has to be covered by at least one locus); (iv) matrix has either at least one complete row or no complete rows. We generated 3,060 matrices that could be analysed with both Terraphast and Gentrius (i.e. with at least one complete row) and 3,060 only with Gentrius (i.e. no complete rows). Additionally we varied minimal amount of missing data per locus (column) and the number of minimally covered species (rows with row sum = 1). For more details on parameters and tested values see Supplementary Figure 3 and Supplementary Table 1. For each of the presence-absence matrix we generated five distinct random trees IQ-TREE 2^21^. The resulting 30,600 simulated instances were analysed with Gentrius to obtain corresponding stands.

#### SIM2

To elucidate the dependence of stand size on the coverage pattern in species and loci we simulated matrices fixing the amount of missing data in loci or species. Namely, we generated 66 matrices with 100 species, 10 and 30 loci, and 30%, 50%, 70% of missing data. First we fixed the number of missing entries per species to uniform (e.g. each species has exactly 3, 5 or 7 missing entries for 10 loci and 30%, 50%, 70% of missing data respectively). The spread across loci varied from uniform (Extended Data Figs. 5A, 6A, matrices with id 1) to datasets with high and low locus coverage (matrices 2-4). Then missing data per loci were fixed and varied for species (Extended Data Figs. 5, 6, matrices 5-11). For all parameter combinations matrices 1-8 did not have species represented by a single locus (i.e. row sum = 1), while matrices 9, 10, 11 had 5, 10, 20 such minimally covered species respectively. All parameters used to generate each matrix are listed in Supplementary Table 3. We next generated 50 distinct random trees with IQ-TREE 2 and computed stands for these trees for each matrix using Gentrius. The same 50 trees were used for each simulated matrix. Note, that we initially simulated multiple matrices per parameter setting. The stand sizes were very similar for similar type of matrices. Hence, the matrices shown in the paper are representative and the conclusions about the dependence of stand sizes on missing data across species and loci are consistent.

### Biological datasets and their phylogenetic trees

We analysed 12 published alignments (Table 1). For each alignment we inferred a maximum likelihood (ML) tree using IQ-TREE 2 (version 2.2.2.4 for D1 and D2, version 2.1.2 for D3-D12) with branch unlinked partition model^9,31^. Best fit evolutionary models for each partition were identified using ModelFinder^32^. For each alignment we computed presence-absence matrix. Next, using Gentrius we generated stands for all 12 inferred ML trees and corresponding matrices. To explore other stands for these 12 empirical presence-absence matrices, for each dataset we also generated 100 distinct random trees using IQ-TREE 2 and computed their stands with Gentrius.

### Topological differences of trees from the same stand

For simulated and empirical datasets D12, D6, D3, D4 we constructed strict consensus trees with IQ-TREE 2. For datasets D8 and D10 with ∼39M and ∼15M trees on corresponding stands we used TreeShredder^33^, which allows efficient analysis for such large tree sets. Importantly, strict consensus trees were only constructed for complete stands. The number of internal branches in strict consensus trees, the tree resolution and the number of multifurcating nodes, as well as their types were computed using a custom R script. The trees were visualised with the same script.

### Data and code availability

The implementation of Gentrius is available in IQ-TREE 2 (since version 2.2, GitHub: https://github.com/iqtree/iqtree2, Website: http://www.iqtree.org/, Gentrius Manual at GitHub: https://github.com/iqtree/iqtree2/tree/master/terrace). The custom matrix simulator is available at GitHub repository: https://github.com/OlgaChern/MatrixSimulator. All simulated and biological data used in this manuscript as well as auxiliary scripts are available at GitHub repository: https://github.com/OlgaChern/Gentrius_2023SM. Note, that all empirical alignments were published previously and are also available via original papers.

## Acknowledgments

We thank Heiko Schmidt and Clement Bader for helpful comments on strict consensus trees. OC and AvH acknowledge funding from the Austrian Science Fund - FWF grant number I-4686.

## Contributions

Conceptualization: OC, CE, AvH

Methodology: OC

Software: OC

Formal Analysis: OC

Investigation: OC

Visualization: OC

Funding acquisition: OC, AvH

Writing – original draft: OC

Writing – review & editing: OC, CE, AvH

## Ethics declarations

### Extended data

Extended Data Figs. 1-6 (provided after Figures and Table of the main text).

## Supplementary information

Supplementary Notes 1-5, Figs. 1–11, and Tables 1–4.

## Extended Data Figures

**Extended Data Fig. 1.**
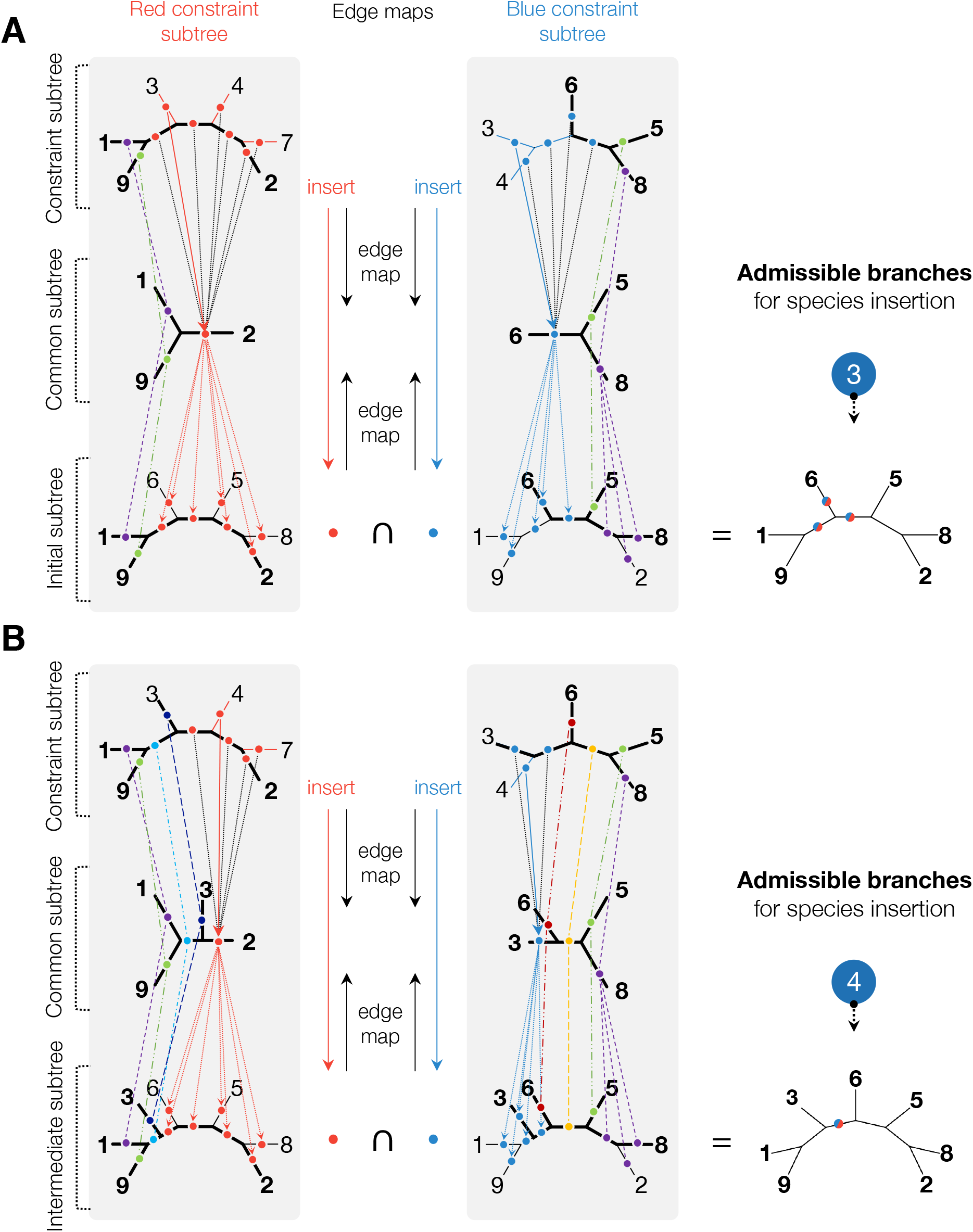
Example of edge maps between pairs of trees and identification of admissible branches. **(A)** The edge maps between initial and constraints subtrees from Fig. 2C before inserting species 3. The edge maps are constructed by collapsing species and branches into branches of the common subtree: from constraint to common subtree and from initial to common subtree. Branches of constraint and initial subtree mapped to the same branch in common subtree are marked by the dots with the same colour. To identify admissible branches for species 3 we look up the maps of its incident branch in constraint subtrees. In red constraint subtree branch incident to species 3 maps to branch incident to species 2 in common subtree, which maps into seven branches marked by red dots in initial subtree. Inserting species 3 on these branches keeps the induced subtree for common species between initial and red constraint subtrees identical. Hence, they are compatible. Similarly, in blue constraint subtree, branch incident to species 3 is mapped to branch incident to species 6, which is mapped to five branches marked by blue dots in initial subtree. The intersection of branches with red and blue dots gives three admissible branches for insertion of species 3. After the insertion each of the three intermediate subtrees is pairwise compatible with red and blue constraint subtrees. **(B)** When species 3 is inserted into one of its admissible branches, the maps have to be updated. Next, in red constraint subtree branch incident to species 4 maps to branch incident to species 2 in common subtree, which is mapped to seven branches with red dots in intermediate subtree. In blue constraint subtree branch incident to species 4 maps to branch incident to species 3 in common subtree, which is mapped to five branches with blue dots in intermediate subtree. The intersection gives only one admissible branch for insertion of species 4. Inserting species 4 on this branch generates an intermediate subtree pairwise compatible with each constraint subtree.

**Extended Data Fig. 2.**
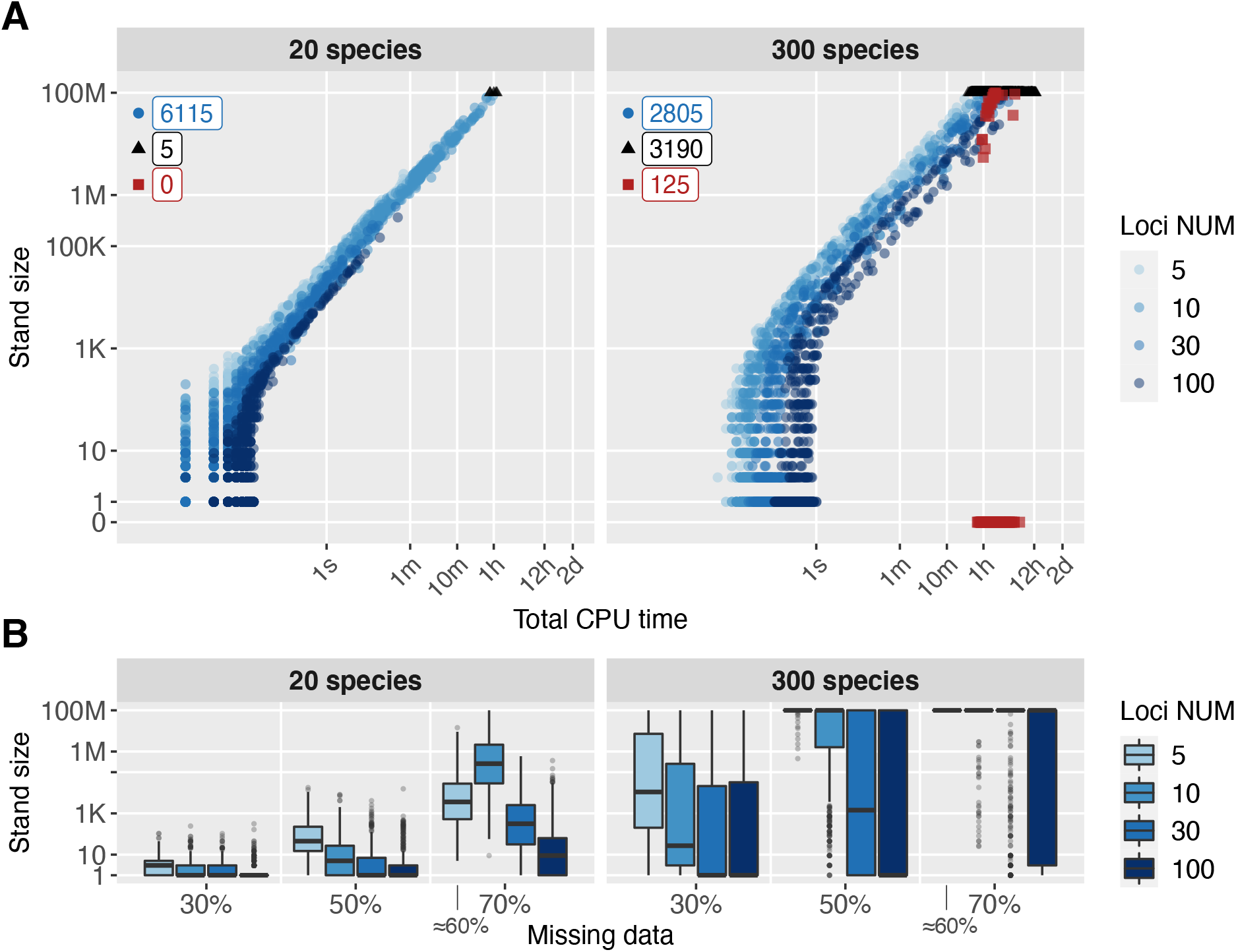
Results of simulation study for datasets with 20 and 300 species. (**A**) Stand size vs. total CPU runtime. Shapes correspond to different computational cases: circles denote complete stands, triangles and squares denote incomplete stands, i.e. triggering MaxStandTrees or MaxIntermediate, respectively. The numbers (top left) specify how often each case occurred. The squares at zero represent complex cases, where MaxIntermediate was triggered, but no tree from a stand was generated yet. For all datasets with red squares, we subsequently ran Gentrius with an alternative initial subtree (Methods), confirming a lower bound of 100M trees on the stand size. (**B**) Influence of the percentage of missing data on stand sizes. Note, that for 5 loci with the constrains imposed (Methods), the resulting presence-absence matrices had around 60% of missing data instead of 70% as required (Supplementary Fig. 4).

**Extended Data Fig. 3.**
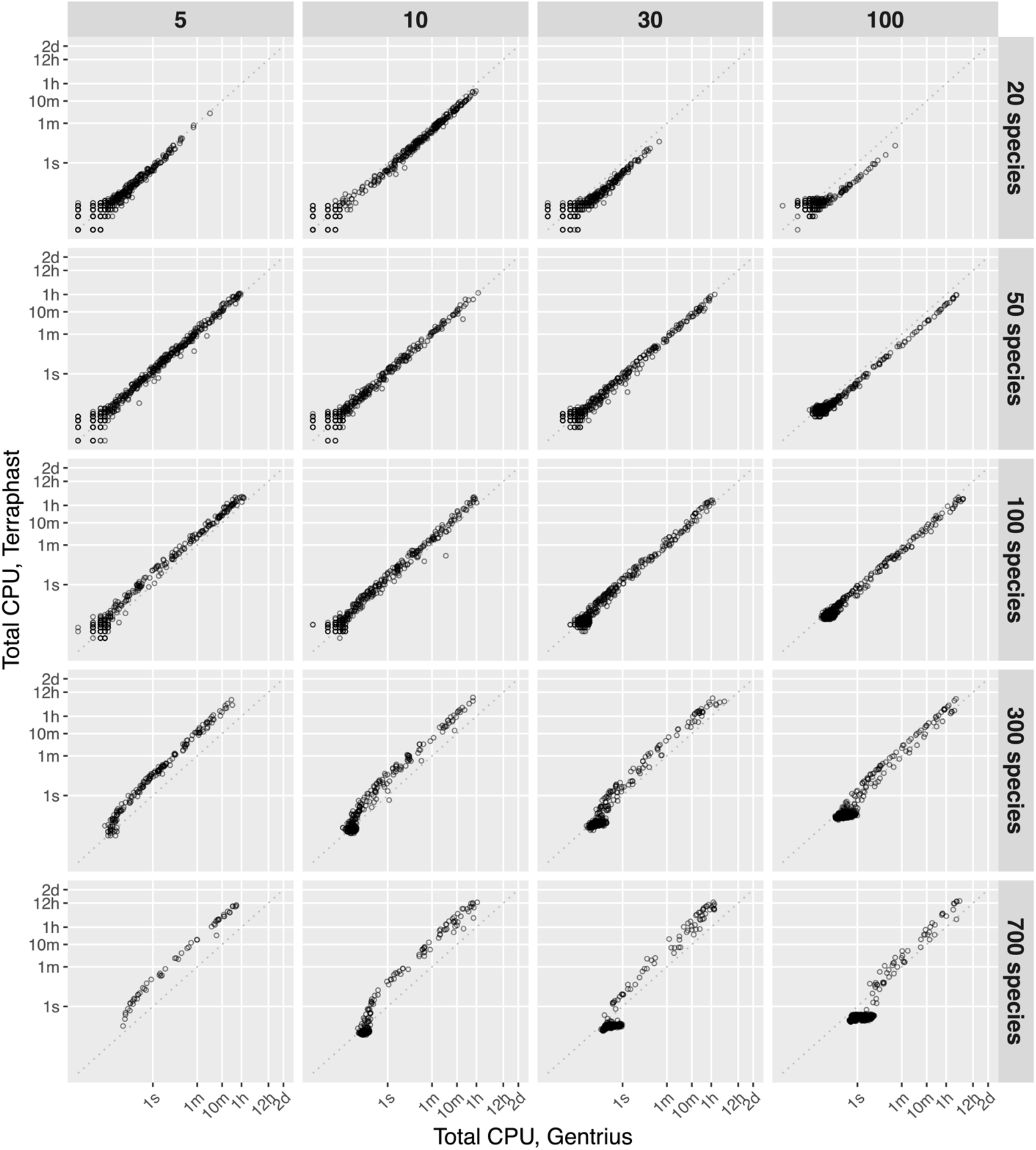
Comparison of total CPU runtimes between Gentrius and Terraphast. The comparison was performed on datasets with at least one comprehensive species (i.e. covered by all loci, corresponds to row without 0’s in the presence-absence species per locus matrix) and with stands less than 100M trees. The plots are arranged by locus number (5 to 100). Gentrius tends to be faster, when species number is larger than locus number. Terraphast is faster, when the locus number is larger than species number. Overall, the runtimes are very comparable.

**Extended Data Fig. 4.**
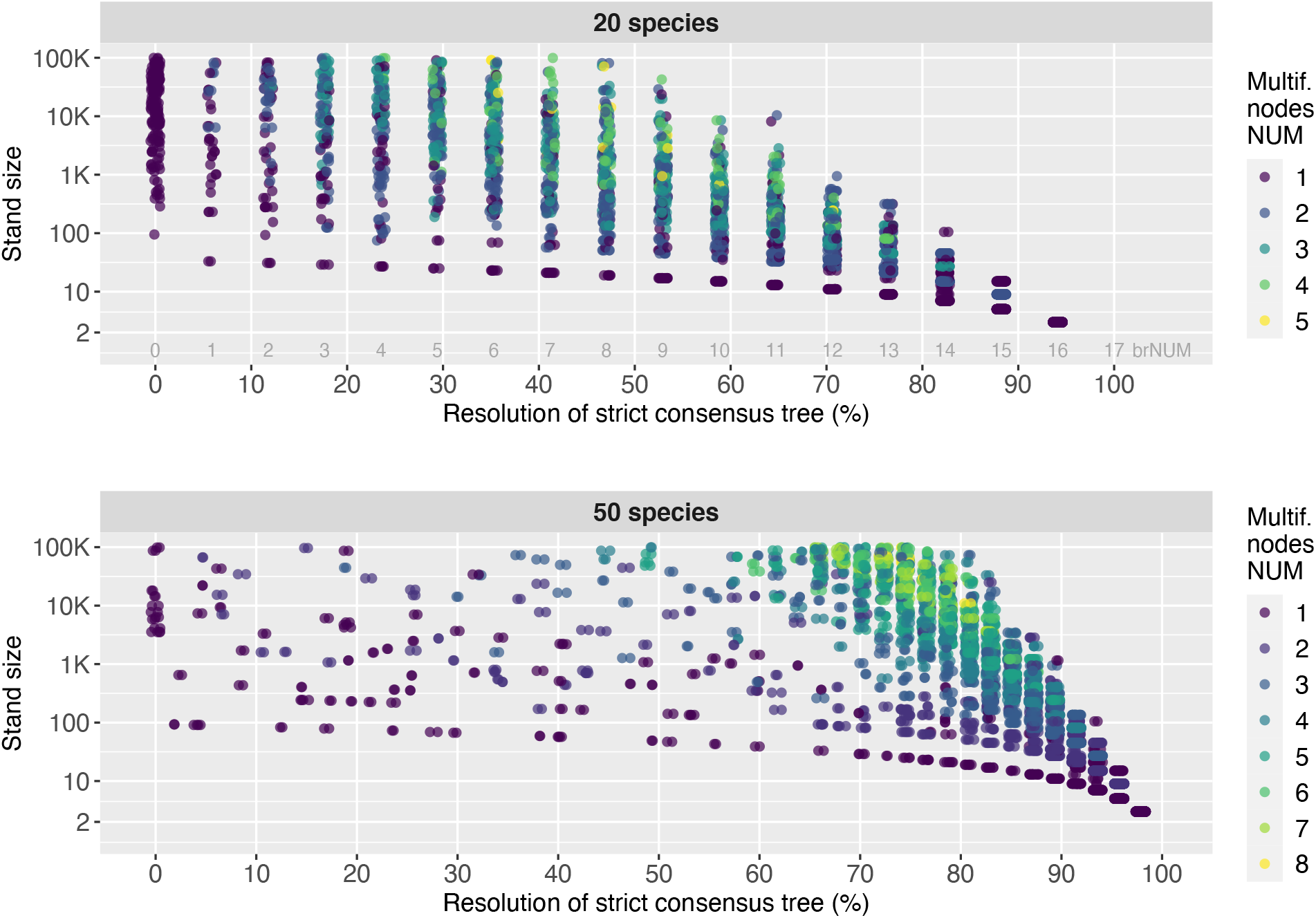
Resolution of strict consensus trees for 20 and 50 species. Each point corresponds to one stand. All stands have up to 100K trees. The resolution refers to the percentage of internal branches on a strict consensus tree compared to the ***n* − 3** internal branches on a fully resolved tree with ***n*** species.

**Extended Data Fig. 5.**
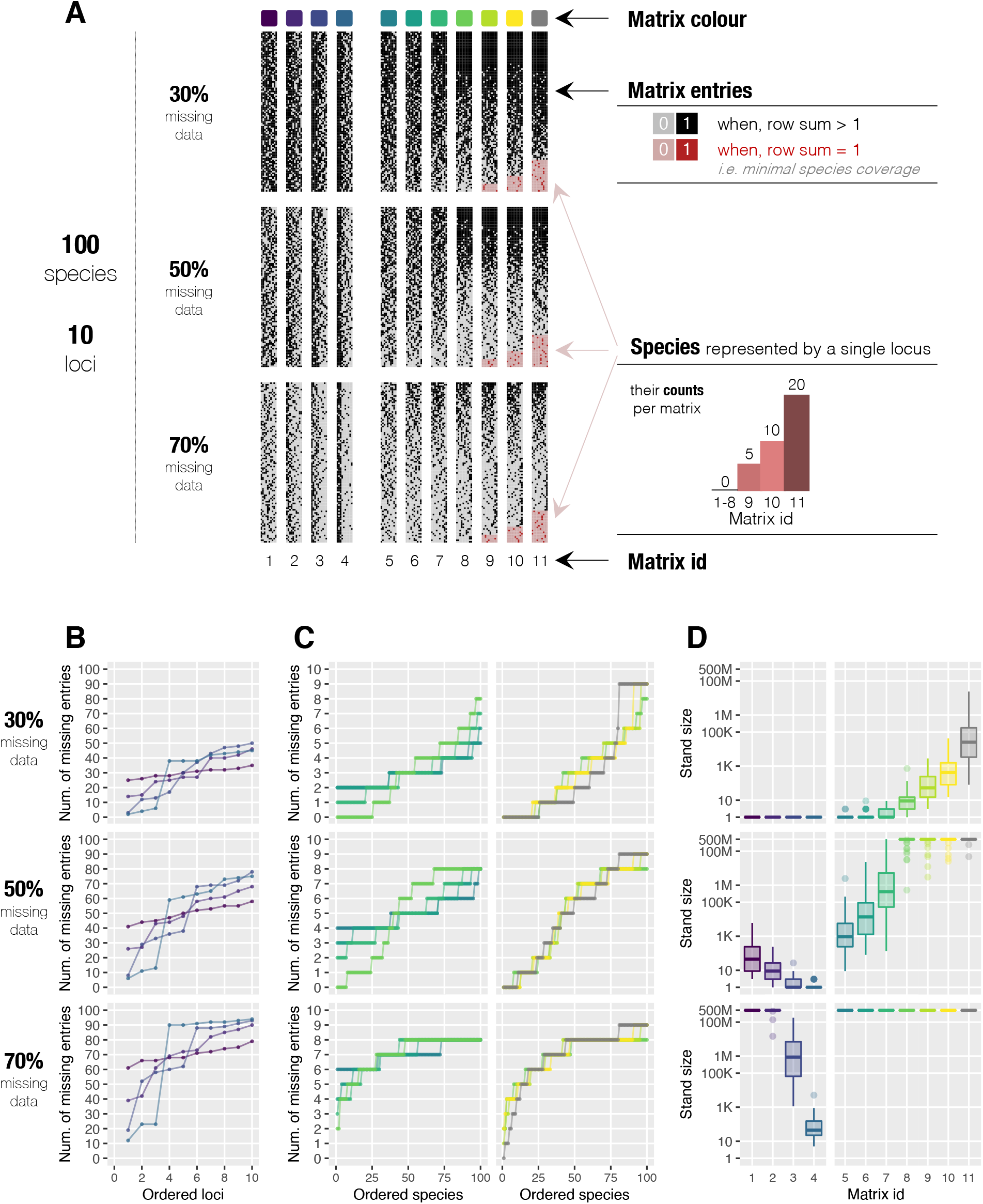
Dependence of stand size on missing data across species and loci. **(A)** The simulated presence-absence matrices have 100 species, 10 loci and 30% to 70% of missing data. Matrices 1 to 4 have fixed number of missing entries per species, namely: 3, 5 and 7 for 30%, 50% and 70% of missing data respectively. Matrices 5 to 11 have fixed number of missing entries per locus. Namely, 30, 50 and 70 for 30%, 50%, and 70% of missing data respectively. **(B)** For matrices 1 to 4 from (A), the summary of missing data across loci. For each matrix the loci are ordered by increasing missing data. The colours correspond to matrix colours in (A). **(C)** For matrices 5 to 11 from (A), the summary of missing data across species. For each matrix the species are ordered by increasing missing data. The colours correspond to matrix colours in (A). **(D)** Stand sizes for matrices from (A). 50 random trees were fixed across all matrices.

**Extended Data Fig. 6.**
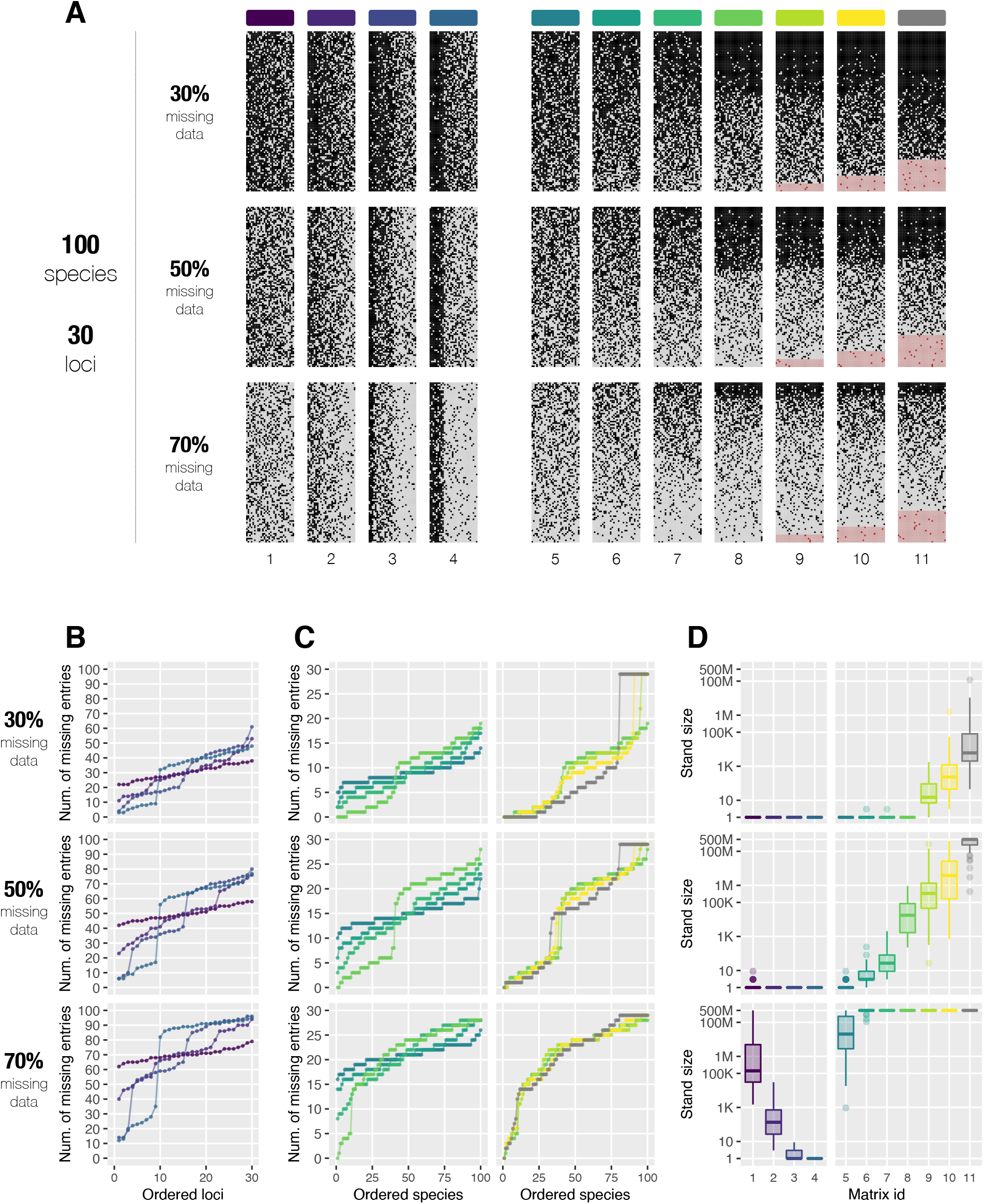
Dependence of stand size on missing data across species and loci. **(A)** The simulated presence-absence matrices have 100 species, 30 loci and 30% to 70% of missing data. Matrices 1 to 4 have fixed number of missing entries per species, namely: 9, 15 and 21 for 30%, 50% and 70% of missing data respectively. Matrices 5 to 11 have fixed number of missing entries per locus. Namely, 30, 50 and 70 for 30%, 50%, and 70% of missing data respectively. **(B)** For matrices 1 to 4 from (A), the summary of missing data across loci. For each matrix the loci are ordered by increasing missing data. The colours correspond to matrix colours in (A). **(C)** For matrices 5 to 11 from (A), the summary of missing data across species. For each matrix the species are ordered by increasing missing data. The colours correspond to matrix colours in (A). **(D)** Stand sizes for matrices from (A). 50 random trees were fixed across all matrices.

