## Supplementary Info for "Gentrius: identifying equally scoring trees in phylogenomics with incomplete data"

**The PDF file includes:**

Supplementary Note 1. Extended description of Gentrius algorithm
Supplementary Note 2. Identifying admissible branches
Supplementary Note 3. Implementation, complexity and validation
Supplementary Note 4. Custom matrix simulator
Supplementary Note 5. Comparison with Terraphast

Supplementary Figs. 1-5
Supplementary Tables 1-4

**Other Supplementary Information for this manuscript includes the following:**

Supplementary Figs. 6-11 (for readability provided as separate files)

#### Supplementary Note 1. Extended description of Gentrius algorithm

1.1. Gentrius – GENerating TRees from Incomplete Unrooted Subtrees
1.2. Tackling computationally complex datasets
1.3. Summary of techniques employed in Gentrius

##### *1.1. Gentrius – GENerating TRees from Incomplete Unrooted Subtrees*

In the following we assume binary unrooted trees. Following definition of (Sanderson et al. 2015), a stand for a set of subtrees is the collection of all trees, which are compatible with these subtrees. Recall, that two trees are called compatible, if there exists a tree, that displays both of them and such a tree exists if and only if two trees have identical induced subtree for their common species (Gordon 1986). For instance, Strict Consensus Merger algorithm (Huson et al. 1999) uses tree compatibility for two trees to sequentially merge a set of incomplete subtrees into a single (multifurcating) strict consensus tree. However, for incomplete subtrees pairwise compatibility does not imply compatibility of the whole set (Song and Hein 2004), i.e. the existence of tree displaying all subtrees is not guaranteed. Therefore, to generate all and only trees from the stand (i.e. a complete stand) all subtrees must be considered simultaneously.

Gentrius generates a stand for a set of (induced) subtrees using the principle of stepwise taxon insertion (also called stepwise addition) under compatibility constraints jointly defined by all subtrees. Both tree compatibility (thus, common subtrees) and stepwise taxon insertion are widely used concepts in phylogenetic methods. For instance, in tree searches for building initial trees (Felsenstein 1993; Olsen et al. 1994; Swofford 2002; Makarenkov and Lapointe 2004; Kozlov et al. 2019; Minh et al. 2020) and for constraint tree search (Minh et al. 2020). Importantly, to the best of our knowledge, none of the previous applications of tree compatibility and stepwise taxon insertion considered multiple constraint subtrees simultaneously, as required for stand generation.

Additionally to the above challenge, stand generation is affected by two computational obstacles: generating even a single tree from a stand is computationally intractable and, moreover, the number of stand trees can be exponential. Together with compatibility and stepwise taxon insertion, Gentrius combines various other techniques and algorithmic advances, crucial to assure generating complete stand in reasonable time.

In the first step, Gentrius selects one of the subtrees to be an initial subtree. Note, that this is the first condition necessary to guarantee generating a complete stand (if feasible due to computational complexity). In Gentrius the initial subtree is chosen to be the subtree with the largest number of species in common with other input subtrees (determined from the presence-absence matrix). When compared to various alternatives, such initial subtrees resulted in generally more efficient stand generation (i.e. shorter run times) as observed in preliminary tests.

Next, species missing from initial subtree are inserted sequentially onto it, while the remaining input subtrees constrain the insertion. Namely, after each insertion of species into a branch, the extended (intermediate) subtree must remain *pairwise compatible* with each constraint subtree.

We call such branches *admissible branches*. Identifying all and only admissible branches is the second condition necessary to generate a complete stand.

To find admissible branches, we developed *edge maps* (Supplementary Note 2), which allow simultaneously tracking compatibility between intermediate subtree and each constraint subtree. Initially, the edge maps are built for each pair of initial and constraint subtrees. After each species insertion the maps have to be updated and admissible branches for a new species are identified with the updated maps. If a species cannot be inserted into an intermediate subtree (i.e. no admissible branches), then Gentrius stops species insertion for this subtree (i.e. encountered a "dead end") and continues with the next intermediate subtree. In fact, Gentrius uses recursion over species and iterates over all admissible branches (i.e. species are inserted and deleted during recursion to generate next intermediate subtree).

A tree from a stand is generated, when all species are inserted. The order of species insertion is determined dynamically and, thus, different stand trees are generated using different species orders. When selecting a new species for insertion Gentrius identifies the species with the smallest number of admissible branches. As confirmed in preliminary tests, the dynamic species order substantially enhances efficiency of Gentrius, resulting in less intermediate trees and dead ends than alternative fixed species orders.

A stand is generated completely, when Gentrius iterated over all admissible branches for all species to be inserted. The whole stand generation process is of branch and bound type.

If feasible in reasonable time, Gentrius generates all and only trees from one stand, i.e. a complete stand (in the worst case impossible due to computational complexity). This is achieved by the choice of initial subtree (must be one of the input subtrees not to miss any stand tree) and by identifying all and only admissible branches for each species. The latter is only possible by simultaneously considering all subtrees thanks to the introduced here edge maps.

The efficiency of Gentrius is attained by the optimal choice of initial subtree (tested among various alternatives), dynamic species order and the corresponding data structure (Supplementary Note 3.3) allowing efficient building and updating of edge maps after insertion and deletion of species during branch and bound like generation of trees.

By combining these various techniques, Gentrius provides to the best of our knowledge the first feasible algorithm for stand generation from unrooted subtrees without any restriction on the input dataset. In the following we describe additional functionality for computationally complex datasets.

##### *1.2. Tackling computationally complex datasets*

Generating even a single tree from a stand is computationally intractable. Therefore, it can happen that Gentrius reaches stopping thresholds without generating any tree. This means that only dead ends were encountered before hitting MaxIntermediate or MaxCPU thresholds (for the description of thresholds see Supplementary Note 3). The default values of thresholds can be modified by the user. However, if there are zero stand trees even for default values, it is already a sign of high complexity of data.

We refer to such datasets as *complex*. In such cases, to exclude from consideration many paths with dead ends, Gentrius constructs an alternative initial subtree. Note, that to generate less dead ends than in the main workflow (i.e. when initial subtree is one of the induced subtrees), the alternative initial subtree must have larger number of species, than any induced subtree. To this end Gentrius constructs alternative initial subtree from the input (complete) tree by removing a user-defined number of species.

However, such an alternative initial subtree comes with a limitation that generating a complete stand is not anymore guaranteed, i.e. Gentrius only provides a lower bound on stand size. Nevertheless, if this lower bound is huge (i.e. excessive number of trees), this is a sufficient sign that missing data hampers phylogenetic inference.

We also provide a script, which uses this alternative approach to find a lower bound specified by the user, imitating MaxStandTrees behaviour. Here, the number of species to be removed from the input tree is chosen dynamically to balance between excluding many dead ends and generating MaxStandTrees trees from the stand.

Thus, enhanced with this alternative initial tree approach, Gentrius can generate complete or partial stands for datasets of all levels of computational difficulty, hence, providing valuable information about multiple equally scoring trees from the same phylogenetic terrace.

##### *1.3. Summary of techniques employed in Gentrius*

Gentrius is the algorithm to generate a complete stand from unrooted subtrees. It combines the following techniques and concepts to assure algorithm correctness and its feasibility

- Stepwise taxon insertion (also called stepwise addition) with constraints (i.e. constraints assure tree compatibility) to recursively generate trees, resulting in branch and bound like strategy for stand generation.
- Tree compatibility is tracked and assured by introduced edge maps: by definition pairwise compatibility is based on shared subtrees for common species; compatibility of the set of (sub)trees is assured by edge maps linking initial/intermediate subtrees with all constraint subtrees. Importantly, all constraint subtrees are considered simultaneously!
- Optimal choice of initial subtree to enhance efficiency.
- Dynamic taxon order to enhance efficiency.
- Alternative initial subtree for complex datasets.

#### Supplementary Note 2. Identifying admissible branches

2.1. Compatibility of trees
2.2. Edge map between a binary unrooted tree and its induced subtree
2.3. Edge map between two binary unrooted trees with overlapping sets of leaves
2.4. Species insertion
2.5. Identifying admissible edges from multiple constraint subtrees
2.6. Final remarks

The correctness of Gentrius, i.e. the guarantee to generate all and only trees from a respective stand, thus, a complete stand, is achieved by two key points: correct choice of initial subtree (must be one of the input (induced) subtrees) and identifying all and only admissible branches for an insertion of missing species. For the latter it is crucial to analyse all subtrees simultaneously.

To determine admissible branches we developed edge maps. These maps are based on the concept of compatibility of two binary unrooted trees and extends to linking intermediate subtree and all constraint trees to allow synchronous analysis. When we want to insert a new species, using edge maps we identify admissible branches for each pair of (initial/intermediate) subtree and constraint subtree. Then the intersection of all "pairwise" admissible branches gives a final list of branches on (initial/intermediate) subtree allowed for the insertion of considered species. The corresponding data structure for building edge maps and their efficient updates during insertion and deletion of species is discussed in Supplementary Note 3.

In the following we provide a formal description of edge maps and prove that they assure identifying all admissible branches. For convenience in mathematical description we use "edge" instead of "branch".

##### *2.1. Compatibility of trees*

Before we introduce edge maps, we first describe the concept of compatibility of trees. Let $T$ be a tree with leaf set (i.e. species) $X$ and $Y \subset X$. The induced subtree $T|_Y$ is obtained from $T$ by removing leaves in $X \backslash Y$ and corresponding branches.

Let $T_1$ and $T_2$ be trees with leaf sets $X_1$ and $X_2$. Trees $T_1$ and $T_2$ are called compatible, if there exists a tree $T$, that displays both of them (Gordon 1986), i.e. $T|_{X_1} = T_1$ and $T|_{X_2} = T_2$.

Let $X_{12} = X_1 \cap X_2$. Trivially, the existence of $T$ requires that $T_1|_{X_{12}} = T_2|_{X_{12}} \coloneqq T_{12}$, i.e. two trees must agree on the relationship for common leaves. If this condition is satisfied, then we can "merge" $T_1$ and $T_2$ through their common subtree $T_{12}$. For instance, such merging can be done by collapsing subtrees for leaves, intrinsic to $T_1$ and $T_2$ (i.e. $X_1 \backslash X_{12}$ and $X_2 \backslash X_{12}$), to branches of $T_{12}$. After merging, these collapsed subtrees are again expanded to obtain tree $T$. In this way, we can obtain one of potentially many trees that are compatible with trees $T_1$ and $T_2$.

The basic idea behind the edge maps for two trees is to keep track of merged parts of two trees (i.e. the induced subtree for common leaves) and the collapsed subtrees special to each of the two considered trees.

##### *2.2. Edge map between a binary unrooted tree and its induced subtree*

Let $T$ be a tree with leaves $X$ and edges $E$. Each edge $e \in E$ is a bipartition of set $X$: $e = A|B$ with $A \cup B = X$ and $A \cap B = \emptyset$. Let $T|_Y$ denote a tree induced by $Y \subset X$ with an edge set $E_Y = \{A \cap Y|B \cap Y: A|B \in E, A \cap Y \neq \emptyset \text{ and } B \cap Y \neq \emptyset\}$. The basic map between edges $E$ and $E_Y$ follows from the definition of an induced tree. Namely, let ε denote the empty edge or no edge, then

$$f': E \to E_Y \cup \{\varepsilon\}$$

$$f'(A|B) = \begin{cases} A \cap Y|B \cap Y, & \text{if } A \cap Y \neq \emptyset \text{ and } B \cap Y \neq \emptyset \\ \varepsilon, & \text{otherwise.} \end{cases}$$

This map was described together with the algorithm to build it in (Chernomor et al. 2016) (therein denoted as map $f$). Here, we will extend the basic map to map every edge from $E$ to some edge in $E_Y$.

Let us consider the following situation at the internal node of $T$ (**Supplementary Fig. 1A**). Here, $A, C$ and $D$ are disjoint sets and $A \cup C \cup D = X$. Further, $A \cap Y = \emptyset$, while $C$ and $D$ have non-empty intersections with $Y$, denoted by $H_1 = C \cap Y$ and $H_2 = D \cap Y$, respectively.

Then $f'(c) = f'(d) = h$, while $a$ and all edges $E_A$ of subtree $T|_A$ map to ε. Since for $a$ both its incident edges $c$ and $d$ map to the same edge $h \in E_Y$, we can unambiguously assign $a$ and all edges in $E_A$ to $h$. Thus, a new map can be defined as

$$f: E \to E_Y,$$

which maps edge $A|B$ to $A \cap Y|B \cap Y$, if intersections are non-empty, and if either intersection is empty the edge is mapped by collapsing corresponding subtree with all $f'(.) = \varepsilon$ to an edge with $f'(.) \neq \varepsilon$ like in the above example.

There can be multiple subtrees with all edges mapped to ε by $f'$, which are collapsed to edges with the same $f'(.) \neq \varepsilon$. **Supplementary Figure 1B** provides a general representation of such a case. Namely, let $C$ and $D$ have non-empty intersections with $Y$ and all leaves between $C$ and $D$ are not in $Y$, i.e. $\forall i$: $A_i \cap Y = \emptyset$. Then it can be easily verified that all edges between $T|_C$ and $T|_D$, denoted by $\mathring{E}_A$, by definition of $f$ are mapped to $C \cap Y|D \cap Y = H_1|H_2 = h$.

In the following we provide conditions for on $C$ and $D$, such that $\mathring{E}_A$ are the only edges, which are mapped to $h$ by $f$.

**Definition:** Let $T$ be a binary unrooted tree with leaf set $X$ and $T|_Y$ is a corresponding induced subtree for some $Y \subset X$. Let for $W \subset X$ the intersections $W \cap Y \neq \emptyset$ and $X \backslash W \cap Y \neq \emptyset$. We call $W$ *irreducible* w.r.t. $T$ and $Y$, if $W$ satisfies one of the following conditions:

(i) $W$ consists of a single leaf: $|W| = 1$; or

(ii) $w_1$ and $w_2$ are two adjacent edges of $w$ in subtree $T|_W$ with leaf sets $W_1$ and $W_2$ (i.e. $W_1 \cup W_2 = W$) and these leaf sets have non-empty intersections with $Y$.

**Supplementary Figure 1C** illustrates the above definition.

**Theorem 1:** Let $T$ be a binary unrooted tree with a leaf set $X$ and edges $E$ as depicted in **Supplementary Figure 1B**. Let $T|_Y$ be a tree induced by some $Y \subset X$ and $E_Y$ denote its edges. Further, let map $f: E \to E_Y$ is defined as above. Then, if $C$ and $D$ are irreducible w.r.t. $T$ and $Y$, then $\mathring{E_A}$ are the only edges, which are mapped to $h$ by $f$.

*Sketch of proof:* The proof is easily obtained using definitions of map $f$ and an irreducible set from above. If $|C| = 1$ (condition (i) of irreducible set w.r.t. $T$ and $Y$), then edge $c$ does not have any adjacent edges. Thus, no additional edge from $T|_C$ maps to $h$. If $C$ satisfies condition (ii), then for edge $c$ and its children edges $c_1$ and $c_2$: $f(c_1) = C_1 \cap Y|D \cap Y$ and $f(c_2) = C_2 \cap Y|D \cap Y$ are different and not equal to $f(c) = C \cap Y|D \cap Y = H_1|H_2 = h$. Note, that since $f(c_1) \neq f(c_2)$, then from the definition of $f$ for all edges in subtrees $T|_{C_1}$ and $T|_{C_2}$ their $f(.) \neq h$. Similar conclusions are obtained for $D$. Thus, $\mathring{E_A}$ are the only edges, which are mapped to $h$.

□

Furthermore, if $|C| = |D| = 1$, i.e. $|X \cap Y| = 2$, then an induced subtree $T|_Y$ consists of a single edge $h = H_1|H_2 = C \cap Y|D \cap Y$ and $f$ maps all edges of $T$ to $h$.

Building edge map $f$ for a tree and its induced subtree takes one tree traversal.

##### *2.3. Edge map between two binary unrooted trees with overlapping sets of leaves*

Let $T_1$ and $T_2$ be trees with leaf sets $Y_1$ and $Y_2$, respectively. Let $Y_{12} = Y_1 \cap Y_2$ denote their common leaves and assume $|Y_{12}| \geq 2$.

If induced trees $T_1|_{Y_{12}} \neq T_2|_{Y_{12}}$, then $T_1$ and $T_2$ are incompatible. Otherwise, let $T_{12}$ denotes their induced subtree for all common leaves, i.e. $T_{12} = T_1|_{Y_{12}} = T_2|_{Y_{12}}$.

We define two maps between edges of trees $T_1$ and $T_2$ and edges of their induced subtree $T_{12}$ as discussed in the previous section

$$f_1: E_1 \to E_{12} \text{ between edges of } T_1 \text{ and } T_{12},$$

$$f_2: E_2 \to E_{12} \text{ between edges of } T_2 \text{ and } T_{12}.$$

Then the map $g: E_1 \to E_2$ between edges of $T_1$ and $T_2$ is defined as

$$g(e) = f_2^{-1} \circ f_1(e) = f_2^{-1}(f_1(e)).$$

**Supplementary Figure 2** illustrates a general case for one edge $h = H_1|H_2$ of $T_{12}$ and all corresponding edges of $T_1$ and $T_2$. Here

$$A_1 \cap Y_{12} = C_1 \cap Y_{12} = H_1 \neq \emptyset$$

$$A_2 \cap Y_{12} = C_2 \cap Y_{12} = H_2 \neq \emptyset$$

Furthermore, let $A_1$ and $A_2$ be irreducible w.r.t. $T_1$ and $Y_{12}$ and let $C_1$ and $C_2$ be irreducible w.r.t. $T_2$ and $Y_{12}$.

If the number of subtrees between $T_1|_{A_1}$ and $T_1|_{A_2}$ (**Supplementary Fig. 2A**) $p > 0$, then

$$\forall i: B_i \cap Y_{12} = \emptyset,$$

i.e. all leaves in $\bigcup_i B_i$ are not present on $T_2$.

Similarly, if the number of subtrees between $T_2|_{C_1}$ and $T_2|_{C_2}$ (**Supplementary Fig. 2C**) $q > 0$, then

$$\forall j: D_j \cap Y_{12} = \emptyset,$$

i.e. all leaves in $\bigcup_j D_j$ are not present on $T_1$.

Denote by $\mathring{E}_B$ all the edges of $T_1$ between subtrees $T_1|_{A_1}$ and $T_1|_{A_2}$ (marked in **Supplementary Fig. 2A** by a dotted box). Then by definition of $f_1$: $\forall e_B \in \mathring{E}_B: f_1(e_B) = h$. Moreover, by theorem 1, $\mathring{E}_B$ are the only edges of $T_1$ that map to $h$.

Similarly, denote by $\mathring{E}_D$ all the edges of $T_2$ between subtrees $T_2|_{C_1}$ and $T_2|_{C_2}$ (marked in **Supplementary Fig. 2C** by a dotted box). Then by definition of $f_2$: $\forall e_D \in \mathring{E}_D: f_2(e_D) = h$. Moreover, by theorem 1, $\mathring{E}_D$ are the only edges of $T_2$ that map to $h$. Then

$$\forall e_B \in \mathring{E}_B: g(e_B) = f_2^{-1} \circ f_1(e_B) = f_2^{-1}\big(f_1(e_B)\big) = f_2^{-1}(h) = \mathring{E}_D$$

and

$$\forall e_D \in \mathring{E}_D: g^{-1}(e_D) = f_1^{-1} \circ f_2(e_D) = f_1^{-1}\big(f_2(e_D)\big) = f_1^{-1}(h) = \mathring{E}_B.$$

##### *2.4. Species insertion*

Let $t \perp e$ denote insertion of leaf (i.e. species) $t$ on edge $e$.

Assume $p > 0$. Then it is trivial to show that for any $t_B \in \bigcup_i B_i$ inserting it on any edge $d \in \mathring{E}_D$, i.e. $t_B \perp d$, generates a new tree $T_2^d$, such that the induce subtrees for common taxa $Y_{12}^* = Y_{12} \cup \{t_B\}$ are identical: $T_1|_{Y_{12}^*} = T_2^d|_{Y_{12}^*} = T_{12}^*$, thus, $T_1$ and newly obtained tree $T_2^d$ are compatible.

Further, inserting $t_B$ on any other edge on $T_2$ not in $\mathring{E}_D$ would lead to different induced subtrees for common leaves, i.e. $T_1|_{Y_{12}^*} \neq T_2^d|_{Y_{12}^*}$.

Therefore, all trees obtained by insertion of leaf $t_B$ on each edge of $\mathring{E}_D$ are the only trees compatible with $T_1$, i.e. tree set $\mathbb{T} = \{T_2^d \big| \, t_B \perp d, \forall d \in \mathring{E}_D\}$.

Since $T_{12}$ and $T_2$ are modified, in order to insert the next leaf on one of such trees $T_2^d$, the maps $f_1$ and $f_2$ have to be updated. Namely, the maps for edges in $\mathring{E}_B$ and $\mathring{E}_D$. Moreover, since all $T_2^d \in \mathbb{T}$ are different, the updated map $f_2^d$ between edges of $T_2^d$ and new induced subtree $T_{12}^*$ are different $\forall d$.

Essentially, for each leaf $t$ from tree $T_1$ not present on tree $T_2$, admissible edges for insertion of this leaf on $T_2$ are $g(e_t)$, where $e_t$ is an incident edge of $t$ in $T_1$. Similarly, for each leaf $t'$ from tree $T_2$ not present on tree $T_1$, its admissible edges on $T_1$ are $g^{-1}(e_{t'})$, where $e_{t'}$ is its incident edge on tree $T_2$.

In summary, w.l.o.g. for each leaf of tree $T_1$ not present on tree $T_2$, map $g$ allows for identification of all admissible edges for insertion of this leaf on $T_2$. Thus, we can generate all trees $\mathbb{T}$, which have leaf $t$ and are compatible with both considered trees. Then one continues with the insertion of other missing leaves on trees from $\mathbb{T}$. Each tree from $\mathbb{T}$ is considered in a pair with $T_1$

(like $T_1$ and $T_2$). The map $g$ is updated for each pair. The next missing leaf is inserted. The process continues and by insertion of all leaves we generate a complete stand for two considered trees.

In the following we describe how to employ such map $g$ to generate stand for many trees.

##### *2.5. Identifying admissible edges from multiple constraint subtrees*

Let $T$ be a tree with leaf set $X$ and edges $E$. Let $T_1, \dots, T_k$ be trees with leaf sets $Y_1, \dots, Y_k$ and edges $E_1, \dots, E_k$, respectively. Denote common leaves between pairs of trees $T$ and $T_i$ by $Y_{Xi} = X \cap Y_i$, $\forall i \in \{1, \dots, k\}$. Then leaves missing on $T$ are $Y = (\bigcup_i Y_i) \backslash (\bigcup_i Y_{Xi})$.

Our aim is to build a stand $S$ for $T$ and all $T_i$ by inserting leaves from $Y$ on $T$. Namely, each tree from $S$ has $X \cup Y$ leaves and is compatible with $T$ and all $T_i$, i.e.

$$\forall T^\circ \in S: T^\circ|_{Y_i} = T_i, \forall i \in \{1, \dots, k\}.$$

For existence of such compatible trees it is necessary (but not sufficient!) that all trees are pairwise compatible. Namely, for each pair of trees their induced subtrees for leaves in common are identical

$$T|_{Y_{Xi}} = T_i|_{Y_{Xi}} \quad (1)$$

$$T_i|_{Y_{ij}} = T_j|_{Y_{ij}} \quad (2)$$

where $Y_{ij} = Y_i \cap Y_j$ and indices $i$ and $j$ run from 1 to $k$.

We are going to insert leaves from $Y$ on $T$ and trees $T_i$ will be used to identifying admissible edges on $T$ for leaf insertion, thus, are termed as constraint trees.

When Eqs. (1) are satisfied, then for each pair of $T_i$ and $T$ we can build maps $g_i: E_i \to E$ between edges of constraint tree $T_i$ and $T$ via their induced subtree for common leaves, i.e. $T_{Xi} = T|_{Y_{Xi}} = T_i|_{Y_{Xi}}$, as discussed in the previous section.

Then for a leaf $t \in Y$ missing from $T$, we identify admissible edges for $t$ based on all constraint trees, which contain $t$

$$E_j^\circ = g_j(e_{tj}),$$

where $j \in J$ and $J$ are indices of constraint trees that contain $t$, $e_{tj}$ are edges incident to $t$ on these constraint trees. Thus, $\forall j \in J$ inserting $t$ on any edge of $E_j^\circ$ on $T$ will generate a new tree compatible with constraint tree $T_j$.

To be compatible with all constraint trees, admissible edges have to be in intersection

$$E_t = \bigcap_j E_j^\circ.$$

If $E_t = \emptyset$, then inserting $t$ on any edge from $E$ on $T$ will violate some constraint tree(s), thus, trees $\{T, T_1, \dots, T_k\}$ are incompatible.

If $E_t \neq \emptyset$, then we iterate over all edges in $E_t$ generating trees on $X \cup \{t\}$ leaves, which are consistent with all $T_i$. After inserting $t$, the maps $g_j: E_j \to E, \forall j \in J$ are updated accordingly.

##### *2.6. Final remarks*

Edge maps allow identifying all admissible branches for a species to be inserted at the current step. Therefore, when all species are inserted, the resulting tree set contains only trees, which are compatible with the input set of subtrees, and also does not miss any tree compatible with these subtrees. Thus, Gentrius generates a complete stand.

**Supplementary Note 3. Implementation, complexity and validation.**

3.1. Inputs
3.2. Data structure
3.3. Complexity
3.4. Stopping thresholds
3.5. Validation

##### *3.1. Inputs*

Gentrius was implemented in IQ-TREE 2 (released in version 2.2) using C++. The program accepts three alternative inputs: either

(i) a tree and species per locus presence-absence matrix (the subtrees are then obtained for each locus by constructing induced trees from the input tree using information from the columns of the matrix); or

(ii) a tree, multi-locus multiple sequence alignment (MSA; i.e. per locus MSAs are concatenated) and information about partitioning of the loci, which are used to obtain induced subtrees; or

(iii) a set of subtrees.

##### *3.2. Data structure*

We used the basic structure of IQ-TREE 2 for trees and build on top of it. A stand data structure consists of

- $T$ - a tree on complete set of taxa (input tree, can be empty, if the input is a set of subtrees);
- $A$ - a presence-absence matrix (input, or derived from the input subtrees);
- $S = \{T|_{a_1}, \dots, T|_{a_k}\}$ - a set of induced (constraint) subtrees with edge sets $E_1, \dots, E_k$, where $a_j$ is the $j$th column of $A$; $S$ is either obtained from $T$ and $A$ or initialised by input subtrees;
- $T'$ - an "agile" tree with edge set $E$, initiated by the initial subtree, which is then modified by insertion and deletion of species to generate stand trees in a recursive manner (by default $T' \in S$, one of the requirement for the correctness of the algorithm), i.e. all and only stand trees are generated;
- $A'$ - a submatrix of presence-absence matrix restricted to species of an agile tree;
- $S' = \{T|_{a_1'}, \dots, T|_{a_k'}\}$ - a set of common subtrees between agile tree and constraint subtrees with edge sets $E_{a_1'}, \dots, E_{a_k'}$;
- $EM$ - the edge maps, which link branches of constraint subtrees to common subtrees, and branches of common subtrees are linked with branches of an agile tree

$$EM = \begin{cases} g_1: E \rightarrow E_{a'_1} \\ \dots \\ g_k: E \rightarrow E_{a'_k} \\ g_1^{-1}: E_{a'_1} \rightarrow E \\ \dots \\ g_k^{-1}: E_{a'_k} \rightarrow E \\ f_1: E_1 \rightarrow E_{a'_1} \\ \dots \\ f_k: E_k \rightarrow E_{a'_k} \\ f_1^{-1}: E_{a'_1} \rightarrow E_1 \\ \dots \\ f_k^{-1}: E_{a'_k} \rightarrow E_k \end{cases}$$

After each species insertion and deletion a submatrix $A'$, an agile tree $T'$, common subtrees $S'$ and edge maps $EM$ are updated.

##### *3.3. Complexity*

Constructing edge maps for each pair of initial and constraint subtrees takes two tree traversals. Updating maps is done only for affected branches. Hence, update does not require complete tree traversal and depends on two considered trees: the larger the species overlap is, the cheaper map update is. The stand generation is in the worst case exponential.

##### *3.4. Stopping thresholds*

Since generating stand trees is in the worst case exponential, to prevent any undesired behaviour with respect to runtime, Gentrius employs three different stopping rules. The run is stopped either when all trees from the stand are generated or if one of conditions is satisfied:

(C1) MaxStandTrees - for a given dataset, $N_1$ trees were generated (default 1 million (M));

(C2) MaxIntermediate - during generation Gentrius generated $N_2$ intermediate trees (i.e. trees with a species number < $n$; default 10M trees); or

(C3) MaxCPU - the CPU time reached $N_3$ hours (default 168 hours (h)).

These thresholds can be modified by the user via corresponding options.

If none of the stopping rules got triggered, the output is a complete stand. Otherwise, the output is a partial stand (can be empty, if MaxIntermediate or MaxCPU is triggered before a tree from a stand is generated). When input is a set of subtrees (i.e. (iii) from above), then if they are incompatible, a “complete” stand is an empty stand, i.e. there are no trees that display all subtrees. However, if the stopping rule was triggered and the stand is empty, this gives no information about compatibility of the input subtrees.

##### *3.5. Validation*

To validate our implementation of Gentrius, we performed the sanity check using all datasets with 50 species and stands with ≤5K trees (in total 4,238 datasets).

The sanity check consisted of the following steps:

1. The input tree is a part of the stand and, thus, must occur among trees generated by Gentrius.
2. All trees in the stand are different.
3. All trees in the stand must have the same set of induced per-locus subtrees.
4. Additional check for datasets with at least one complete species (row) (2,079 out of 4,238 datasets tested for sanity check): the stand size should be the same between Gentrius and Terraphast.

All tested datasets fulfilled the sanity check.

**Supplementary Note 4. Custom matrix simulator**

To simulate presence-absence of data for species in loci, we developed a custom matrix simulator (GitHub repository: https://github.com/OlgaChern/MatrixSimulator), which simulates a random binary matrices according to a number of different parameters controlling the amount and distribution of zeros (i.e. missing data) in a matrix. The simulation begins with a matrix of 1's. The rows and columns are split into three categories, which are assigned with different user-defined sample probabilities. Then a pair of row and column indices is sampled. If the corresponding matrix entry is 0, a pair of indices is resampled. If the matrix entry equals 1, it is assigned to 0. The process continues until a required number of 0's (i.e. the amount of missing data) is reached or if no more zeros can be allocated without violating constraints on the minimal and maximal numbers of 0's and 1's in each row and column. These constraints can lead to a smaller number of zeros allocated at the boundary of parameters.

To reduce the number of row/column sampling categories one can simply set the size of category to 0 via input parameters. The simulator can be also used to generate more complex matrices by combining multiple matrices generated using different parameter values.

The following parameters were employed in our simulations: (P1) minimal number of 0's in each row; (P2) number of rows with minimal coverage (row sum = 1); (P3) minimal number of 0's in each column; (P4) different distributions of 0's across rows (i.e. species are covered by different number of loci); (P5) different distributions of 0's across columns (i.e. loci cover different number of species). All these parameters control the distribution of 0's (i.e. missing data) in a species per locus presence-absence matrix for the triplet comprised of the number of species, loci and the percentage of missing data. For other available parameters refer to the simulator manual.

Note, if minimum % of zeros per column equals the % of missing data in a matrix, then trivially the distribution of zeros across columns is uniform. Namely, the simulator iterates over columns and assigns zeros using sample probabilities of rows. Since no more additional zeros have to be allocated, simulation stops.

#### Supplementary Note 5. Comparison with Terraphast

Terraphast is based on the algorithm developed for rooted trees. Thus, it can only analyse datasets that have at least one species with no missing data, which is treated as a root.

By design exactly half (3,060) of our simulated matrices contained at least one complete species (row) and, thus, could be analysed by Terraphast and Gentrius. However, Terraphast does not have any stopping rules. Therefore, to be able to compare runtimes between two software and to prevent any runtime issues, we applied Terraphast only on datasets, which had stands with less than $100M$ trees, as computed by Gentrius (in total 10,293 datasets).

Moreover, for comparison of CPU runtimes, we do not require trees. Therefore, we modified the code of Terraphast, by adding an option to turn off writing generated trees into a file. This required adding three lines in the file `terraphast/terraphastI/app/main.cpp` marked below by the black box:

```
#include <bitset>
#include <chrono>
#include <fstream>
#include <iomanip>
#include <iostream>
#include <sstream>
#include <thread>

#include <terraces/simple.hpp>

int main(int argc, char** argv) try {
        auto tree_file_name = std::string{};
        auto data_file_name = std::string{};
        if (argc == 2) {
                tree_file_name = argv[1] + std::string{".nwk"};
                data_file_name = argv[1] + std::string{".data"};
        } else if (argc == 3) {
                tree_file_name = argv[1];
                data_file_name = argv[2];
        } else {
                std::cerr << "Usage: \n"
                          << argv[0] << " <tree-file> <occurrence file>\n"
                          << argv[0] << " <common-basename>\n";
                return 1;
        }
        auto trees = std::ostringstream{};
        const auto terraces_count =
                terraces::simple::print_terrace_from_file(tree_file_name, data_file_name, trees);

        std::cout << "There are " << terraces_count
                  << " trees on the terrace.\n\nThe trees in question are:\n"
                  << trees.str() << '\n';
} catch (std::exception& e) {
        std::cerr << "Error: " << e.what() << "\n";
}
```

### Supplementary Figures and Tables

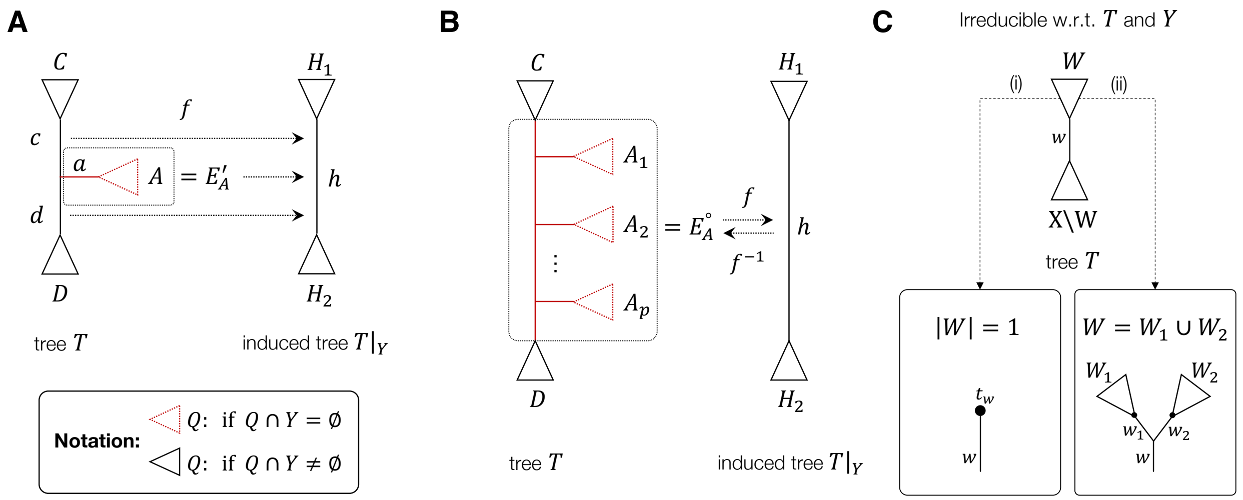


**Supplementary Figure 1 | Illustrations of combinatorial correspondence between a tree and its induced subtree.** Triangles indicated subtrees. Black solid and red dotted triangles indicate that corresponding leaf set has a non-empty or empty overlap with $Y$, respectively. (A) Tree $T$ and its induced subtree $T|_Y$ for leaf set $Y$. A local case at the internal node with $C \cap Y \neq \emptyset$, $D \cap Y \neq \emptyset$, $A \cap Y = \emptyset$. Edges $E'_A$ are all red edges within dotted box, including edges of a subtree marked by red triangle. Map $f$ assigns all edges in $E'_A$ to $h$. (B) Tree $T$ and its induced subtree $T|_Y$ for leaf set $Y$. General case. Here, $C \cap Y \neq \emptyset$ and $D \cap Y \neq \emptyset$ and both $C$ and $D$ are irreducible w.r.t. $T$ and $Y$. Further, $\forall i\colon A_i \cap Y = \emptyset$. Edges $E^{\circ}_A$ are all edges between subtrees corresponding to $C$ and $D$ (red edges within dotted box, including edges of subtrees marked by red triangles). From theorem 1 $E^{\circ}_A$ are the only edges, which map to $h$ via $f$. Namely, $\forall e \in E^{\circ}_A\colon f(e) = h$ and $\forall e' \notin E^{\circ}_A\colon f(e') \neq h$. (C) Illustration for definition of leaf set $W$ irreducible w.r.t. $T$ and $Y$. Two possible topologies for subtree with leaf set $W$.

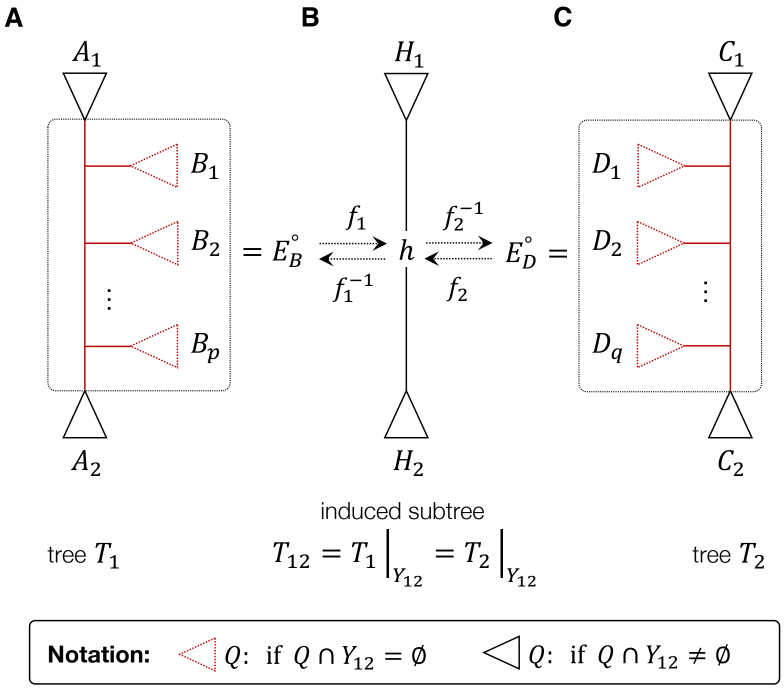


**Supplementary Figure 2 | Edge maps between two trees $T_1$ and $T_2$ via induced subtree for their common species $Y_{12}$.** Triangles indicated subtrees. Black solid and red dotted triangles indicate that corresponding leaf set has a non-empty or empty overlap with $Y_{12}$, respectively. (A) Tree $T_1$ with $A_1 \cap Y_{12} = H_1 \neq \emptyset$, $A_2 \cap Y_{12} = H_2 \neq \emptyset$ and both $A_1$ and $A_2$ are irreducible w.r.t. $T_1$ and $Y_{12}$. Further, $\forall i: B_i \cap Y_{12} = \emptyset$. Edges $E_B^\circ$ are all edges between subtrees corresponding to $A_1$ and $A_2$ (red edges within dotted box, including edges of subtrees marked by red triangles). $E_B^\circ$ are the only ones, which map to $h$ via $f_1$. Namely, $\forall e \in E_B^\circ: f_1(e) = h$ and $\forall e' \notin E_B^\circ: f_1(e') \neq h$. (B) Induced subtree of trees $T_1$ and $T_2$ for their common taxa $Y_{12}$. (C) Tree $T_2$ with $C_1 \cap Y_{12} = H_1 \neq \emptyset$, $C_2 \cap Y_{12} = H_2 \neq \emptyset$ and both $C_1$ and $C_2$ are irreducible w.r.t. $T_2$ and $Y_{12}$. Further, $\forall j: D_j \cap Y_{12} = \emptyset$. Edges $E_D^\circ$ are all edges between subtrees corresponding to $D_1$ and $D_2$ (red edges within dotted box, including edges of subtrees marked by red triangles). $E_D^\circ$ are the only ones, which map to $h$ via $f_2$. Namely, $\forall e \in E_D^\circ: f_2(e) = h$ and $\forall e' \notin E_D^\circ: f_2(e') \neq h$. Inserting any leaf $t_B$ from $\cup_i B_i$ to any of the edge in $E_D^\circ$ keeps the induced subtree for common species between $T_1$ and modified $T_2$ identical. Thus, after insertion of $t_B$ trees remain compatible.

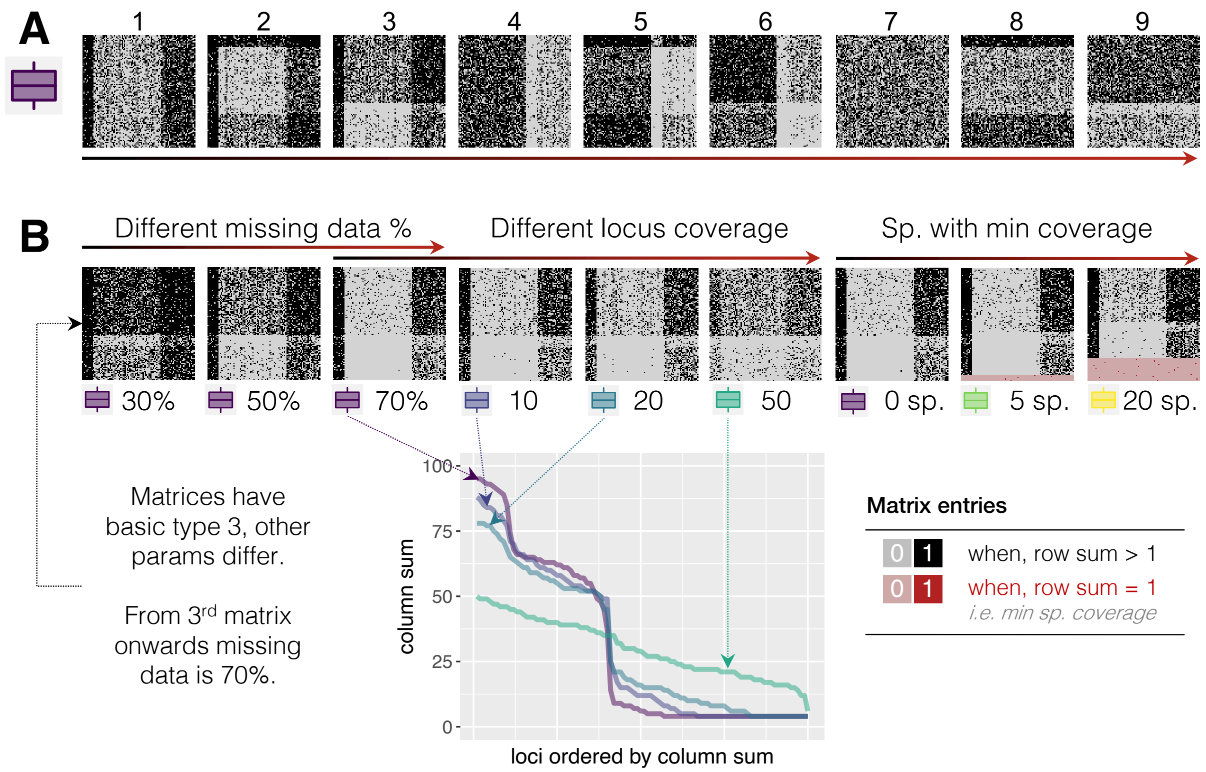


**Supplementary Figure 3 | Examples of different distributions of missing data across matrices (SIM1). (A)** The nine basic types exemplified on matrices for 100 species, 100 loci and 50% of missing data. Here, each row and each column has at least one zero (i.e. missing data). These types are based on combinations of rows and columns with varying amounts of missing data (low, medium, high, or uniform, see also Supplementary Table 1). Missing entries (zeros) are indicated by grey dots. The black-red arrow points in the direction of difficulty (i.e. in general, increased stand sizes). For basic types the minimum number of zeros per locus is one and the number of species covered by a single locus (i.e. minimal coverage, row sum = 1) is null. **(B)** Examples of matrices with basic type 3 and varying other parameters: the percentage of missing data, locus coverage and the number of species with minimal coverage. The locus coverage refers to the column sums. The parameter for locus coverage is the minimal number of missing entries per locus (i.e. per column). The graphic shows locus coverage by means of column sums for four matrices with the same amount of missing data (70%). The species with minimal coverage are species with a non-missing entry in a single locus (i.e. row sum = 1). The black-red arrows point in the direction of difficulty (i.e. in general, increased stand sizes).

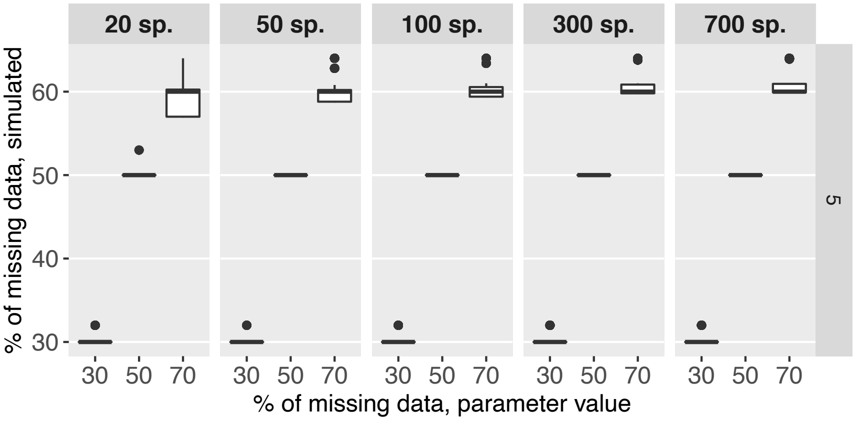


**Supplementary Figure 4 | Comparison of missing data in the presence-absence matrices parameter vs. simulated for five loci.** For five loci the % of missing data in the simulated matrices slightly diverges from the parameter used to run the simulator. It is larger for some simulated matrices, which contain species with minimal coverage (e.g. for 20 species, 4 species with minimal coverage require 16 zeros + 16 zeros for each other row for the setting with at least one missing value in each row, thus, a minimum of 32% of missing data is required in the simulated matrix to fulfil the constraint imposed on the matrix). The % of missing data in the simulated matrix is smaller, if increasing missing entries violates the constraints on column and row sums (e.g. simulated matrices cannot have 70% without violating row/column sum constraints; therefore, the simulated matrices have approximately 60% of missing data). Note, that in our simulations, for all other locus number values the % of missing data in the simulated matrix equals to the used parameter (i.e. 30%, 50%, 70%). Therefore, they are not shown in this plot.

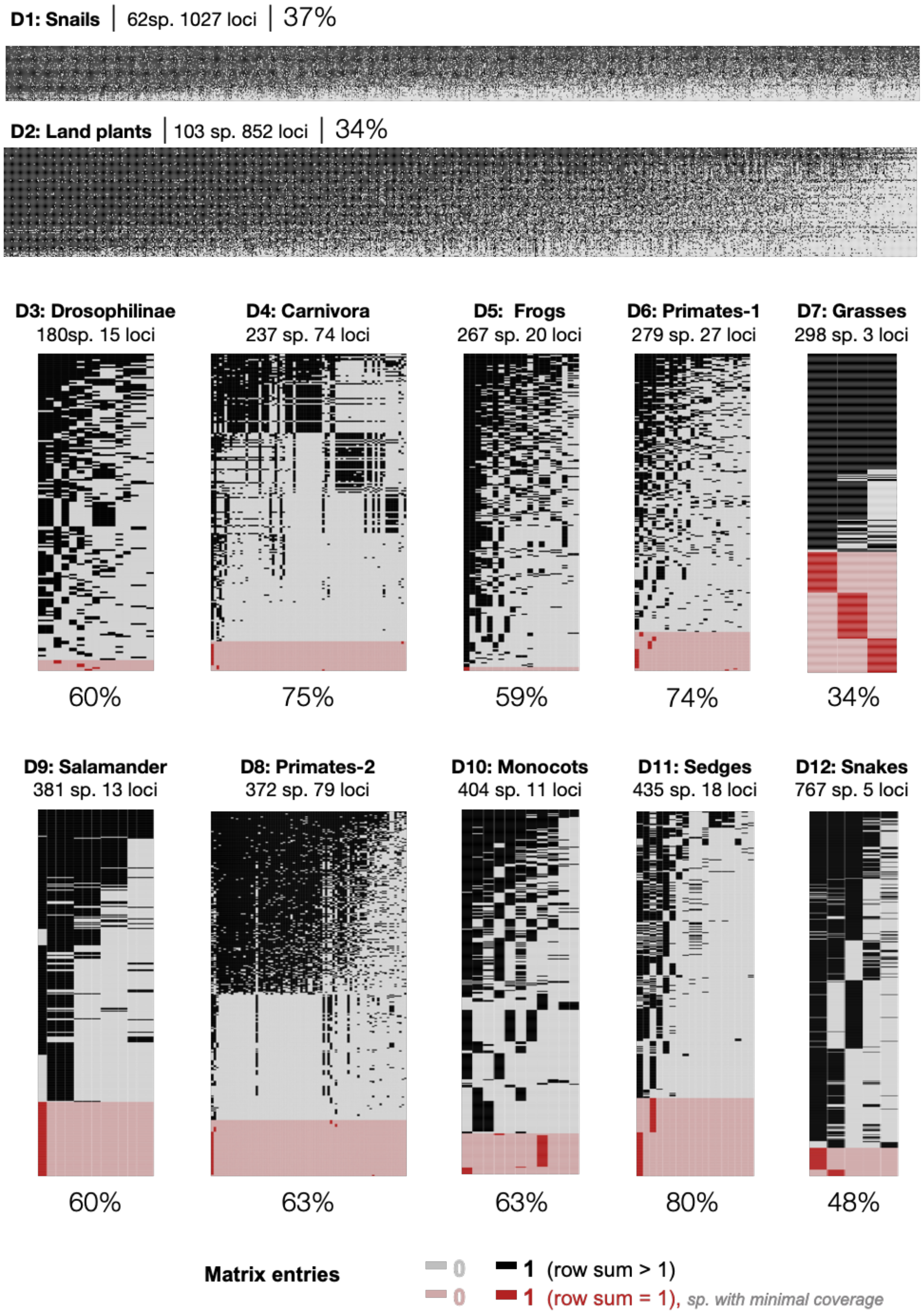


**Supplementary Figure 5 | Visualisation of species per locus presence-absence matrices for biological datasets in Table 1 (main text).** The percentages below each matrix indicate the percentage of missing data in the corresponding matrix (i.e. grey and pale red entries, which stand for "0" in the matrix). Rows in red correspond to species with minimal coverage, i.e. only one non-missing entry (row sum = 1).

**Supplementary Figure 6 (separate file)**

**Strict consensus tree of trees from a stand for ML tree inferred for Snakes (D12).** Provided in a separate file to allow for high resolution. S – stand size, R – tree resolution, iB – number of internal branches, mN – number of multifurcating nodes.

**Supplementary Figure 7 (separate file)**

**Strict consensus tree of trees from a stand for ML tree inferred for Primates-1 (D7).** Provided in a separate file to allow for high resolution. S – stand size, R – tree resolution, iB – number of internal branches, mN – number of multifurcating nodes.

**Supplementary Figure 8 (separate file)**

**Strict consensus tree of trees from a stand for ML tree inferred for Drosophilinae (D3).** Provided in a separate file to allow for high resolution. S – stand size, R – tree resolution, iB – number of internal branches, mN – number of multifurcating nodes.

**Supplementary Figure 9 (separate file)**

**Strict consensus tree of trees from a stand for ML tree inferred for Primates-2 (D8).** Provided in a separate file to allow for high resolution. S – stand size, R – tree resolution, iB – number of internal branches, mN – number of multifurcating nodes.

**Supplementary Figure 10 (separate file)**

**Strict consensus tree of trees from a stand for ML tree inferred for Monocots (D10).** Provided in a separate file to allow for high resolution. S – stand size, R – tree resolution, iB – number of internal branches, mN – number of multifurcating nodes.

**Supplementary Figure 11 (separate file)**

**Strict consensus tree of trees from a stand for ML tree inferred for Carnivora (D4).** Provided in a separate file to allow for high resolution. S – stand size, R – tree resolution, iB – number of internal branches, mN – number of multifurcating nodes.

**Supplementary Table 1: Parameters employed for generating presence-absence species per locus matrices using custom matrix simulator.** Combination of rows/columns with different spread of missing data resulted in nine base types of matrices for a triple of (n, k, m). Minimum number of 0's in each row (r0) is used to generate matrices with and without complete rows. The number of species with minimal coverage (u) was varied only when parameter on the minimum number of 0's in each column c0 = 1. For other c0 values u = **0**. Further, c0 = 50% can only be used with m ≥ 50%, since it requires at least 50% of missing entries in the matrix.

| | | | | |
|---|---|---|---|---|
| • **Matrix size** | | | | |
| Number of species (rows) | **n** | 20, 50, 100, 300, 700 | | |
| Number of loci (columns) | **k** | 5, 10, 30, 100 | | |
| • **Missing data (0's) in the matrix** | | | | |
| **Percentage of 0's** | **m** | 30%, 50%, 70% | | |
| **Distribution of 0's** | | | | |
| *Types of row/col categories* | | *Type 1* | *Type 2* | *Type 3* |
| % of rows/cols in categories out of **n-u** rows/ **k** cols | **(rf1, rf2, rf3)** **(cf1, cf2, cf3)** | (100, 0, 0) | (60, 10, 30) | (10, 60, 30) |
| Sampling probabilities of rows/cols in categories | **(rp1, rp2, rp3)** **(cp1, cp2, cp3)** | (1, 0, 0) | (0.1, 0.6, 0.3) | (0.05, 0.7, 0.25) |
| The main feature of the type | | uniform distribution of missing entries in rows/cols | small number of rows/cols with large amounts of missing entries | small number of rows/cols with high coverage (small amount of missing entries) |
| *Additional parameters* | | | | |
| Minimum number of 0's in each row | **r0** | null (no constraint), and at least one row is complete (all 1's)* | at least one 0 in each row | |
| Minimum number (or %) of 0's in each column | **c0** | 1, 10%, 20%, 50% | | |
| % of rows with minimal coverage (row sum = 1) | **u** | 0, 5%, 20% | | |
| distribution of species with minimal coverage across columns from different categories | | uniformly random<br>Note: All values in corresponding row are set to 0's. Then one column is sampled uniformly at random and an entry is set to 1. | | |

* datasets with at least one complete row can be analysed also with Terraphast

**Supplementary Table 2: Summary of CPU runtime of Gentrius for simulated datasets.** Number of datasets whose run finished within corresponding time range. Here, s, m, h, d denote seconds, minutes, hours, and days, respectively. MaxStandTrees and MaxIntermediate were set to 100M each.

| **Taxa** | <= 1s | (1s, 1m] | (1m, 30m] | (30m, 1h] | (1h, 2h] | (2h, 12h] | (12h, 1d] | (1d, 1.5d] |
|---|---|---|---|---|---|---|---|---|
| 20 | 5,517 | 447 | 145 | 8 | 3 | 0 | 0 | 0 |
| 50 | 4,480 | 579 | 278 | 265 | 459 | 59 | 0 | 0 |
| 100 | 3,407 | 477 | 331 | 631 | 883 | 391 | 0 | 0 |
| 300 | 2,282 | 300 | 178 | 932 | 1,304 | 1,115 | 9 | 0 |
| 700 | 1,155 | 721 | 175 | 1,080 | 1,482 | 1,260 | 219 | 28 |

**Supplementary Table 3: Parameters employed for generating presence-absence species per locus matrices using custom matrix simulator for SIM2.** For parameter meaning refer to Supplementary Table 1. Here, all matrices have 100 species (-n 100). The values in grey (triplets [-r0, -rf, -rp], or [-c0, -cf -cp]) set uniform sampling probabilities for rows or columns (i.e. uniform distribution of zeros across rows/species or columns/loci). Other parameters in black were chosen and tuned to provide the variation of spread of missing values across rows/columns (species/loci). The values in pale pink indicate absence of minimally covered species (i.e. row sum = 1).

| Loci (-k) | % of missing data (-m) | Matrix id | Other parameters | | | | | | |
|---|---|---|---|---|---|---|---|---|---|
| | | | -r0 | -c0 | -u | -rf | -rp | -cf | -cp |
| 10 | 30 | 1 | 3 | 23 | 0 | 0 0 100 | 0 0 1 | 0 0 100 | 0 0 1 |
| | | 2 | 3 | 7 | 0 | 0 0 100 | 0 0 1 | 25 45 30 | 0.15 0.3 0.5 |
| | | 3 | 3 | 2 | 0 | 0 0 100 | 0 0 1 | 10 40 50 | 0.05 0.25 0.7 |
| | | 4 | 3 | 2 | 0 | 0 0 100 | 0 0 1 | 10 20 70 | 0.05 0.1 0.85 |
| | | 5 | 2 | 30 | 0 | 0 0 100 | 0 0 1 | 0 0 100 | 0 0 1 |
| | | 6 | 2 | 30 | 0 | 25 45 30 | 0.15 0.3 0.5 | 0 0 100 | 0 0 1 |
| | | 7 | 1 | 30 | 0 | 10 40 50 | 0.05 0.25 0.7 | 0 0 100 | 0 0 1 |
| | | 8 | 0 | 30 | 0 | 10 30 60 | 0.05 0.1 0.85 | 0 0 100 | 0 0 1 |
| | | 9 | 0 | 30 | 5 | 10 30 60 | 0.05 0.1 0.85 | 0 0 100 | 0 0 1 |
| | | 10 | 0 | 30 | 10 | 10 30 60 | 0.05 0.1 0.85 | 0 0 100 | 0 0 1 |
| | | 11 | 0 | 30 | 20 | 10 30 60 | 0.05 0.1 0.85 | 0 0 100 | 0 0 1 |
| | 50 | 1 | 5 | 40 | 0 | 0 0 100 | 0 0 1 | 0 0 100 | 0 0 1 |
| | | 2 | 5 | 20 | 0 | 0 0 100 | 0 0 1 | 25 45 30 | 0.15 0.3 0.5 |
| | | 3 | 5 | 5 | 0 | 0 0 100 | 0 0 1 | 10 40 50 | 0.05 0.25 0.7 |
| | | 4 | 5 | 5 | 0 | 0 0 100 | 0 0 1 | 10 20 70 | 0.05 0.1 0.85 |
| | | 5 | 4 | 50 | 0 | 0 0 100 | 0 0 1 | 0 0 100 | 0 0 1 |
| | | 6 | 3 | 50 | 0 | 25 45 30 | 0.15 0.3 0.5 | 0 0 100 | 0 0 1 |
| | | 7 | 2 | 50 | 0 | 10 40 50 | 0.05 0.25 0.7 | 0 0 100 | 0 0 1 |
| | | 8 | 0 | 50 | 0 | 10 30 60 | 0.05 0.1 0.85 | 0 0 100 | 0 0 1 |
| | | 9 | 0 | 50 | 5 | 10 30 60 | 0.05 0.1 0.85 | 0 0 100 | 0 0 1 |
| | | 10 | 0 | 50 | 10 | 10 30 60 | 0.05 0.1 0.85 | 0 0 100 | 0 0 1 |
| | | 11 | 0 | 50 | 20 | 10 30 60 | 0.05 0.1 0.85 | 0 0 100 | 0 0 1 |
| | 70 | 1 | 7 | 60 | 0 | 0 0 100 | 0 0 1 | 0 0 100 | 0 0 1 |
| | | 2 | 7 | 35 | 0 | 0 0 100 | 0 0 1 | 25 45 30 | 0.15 0.3 0.5 |
| | | 3 | 7 | 5 | 0 | 0 0 100 | 0 0 1 | 10 40 50 | 0.05 0.25 0.7 |
| | | 4 | 7 | 5 | 0 | 0 0 100 | 0 0 1 | 10 20 70 | 0.05 0.1 0.85 |
| | | 5 | 6 | 70 | 0 | 0 0 100 | 0 0 1 | 0 0 100 | 0 0 1 |
| | | 6 | 4 | 70 | 0 | 25 45 30 | 0.15 0.3 0.5 | 0 0 100 | 0 0 1 |
| | | 7 | 3 | 70 | 0 | 10 40 50 | 0.05 0.25 0.7 | 0 0 100 | 0 0 1 |
| | | 8 | 1 | 70 | 0 | 10 20 70 | 0.1 0.25 0.65 | 0 0 100 | 0 0 1 |
| | | 9 | 1 | 70 | 5 | 10 20 70 | 0.1 0.25 0.65 | 0 0 100 | 0 0 1 |
| | | 10 | 1 | 70 | 10 | 10 30 60 | 0.1 0.25 0.65 | 0 0 100 | 0 0 1 |
| | | 11 | 0 | 70 | 20 | 10 30 60 | 0.05 0.3 0.65 | 0 0 100 | 0 0 1 |
| 30 | 30 | 1 | 9 | 20 | 0 | 0 0 100 | 0 0 1 | 0 0 100 | 0 0 1 |
| | | 2 | 9 | 7 | 0 | 0 0 100 | 0 0 1 | 25 45 30 | 0.15 0.3 0.5 |
| | | 3 | 9 | 2 | 0 | 0 0 100 | 0 0 1 | 10 40 50 | 0.05 0.25 0.7 |
| | | 4 | 9 | 2 | 0 | 0 0 100 | 0 0 1 | 10 20 70 | 0.05 0.1 0.85 |
| | | 5 | 5 | 30 | 0 | 0 0 100 | 0 0 1 | 0 0 100 | 0 0 1 |
| | | 6 | 2 | 30 | 0 | 25 45 30 | 0.15 0.3 0.5 | 0 0 100 | 0 0 1 |
| | | 7 | 2 | 30 | 0 | 10 40 50 | 0.05 0.25 0.7 | 0 0 100 | 0 0 1 |
| | | 8 | 0 | 30 | 0 | 10 30 60 | 0.05 0.1 0.85 | 0 0 100 | 0 0 1 |
| | | 9 | 0 | 30 | 5 | 10 30 60 | 0.05 0.1 0.85 | 0 0 100 | 0 0 1 |
| | | 10 | 0 | 30 | 10 | 10 30 60 | 0.05 0.1 0.85 | 0 0 100 | 0 0 1 |
| | | 11 | 0 | 30 | 20 | 10 30 60 | 0.05 0.1 0.85 | 0 0 100 | 0 0 1 |
| | 50 | 1 | 15 | 40 | 0 | 0 0 100 | 0 0 1 | 0 0 100 | 0 0 1 |
| | | 2 | 15 | 20 | 0 | 0 0 100 | 0 0 1 | 25 45 30 | 0.15 0.3 0.5 |
| | | 3 | 15 | 5 | 0 | 0 0 100 | 0 0 1 | 10 40 50 | 0.05 0.25 0.7 |
| | | 4 | 15 | 5 | 0 | 0 0 100 | 0 0 1 | 10 20 70 | 0.05 0.1 0.85 |
| | | 5 | 10 | 50 | 0 | 0 0 100 | 0 0 1 | 0 0 100 | 0 0 1 |
| | | 6 | 5 | 50 | 0 | 25 45 30 | 0.15 0.3 0.5 | 0 0 100 | 0 0 1 |
| | | 7 | 3 | 50 | 0 | 10 40 50 | 0.05 0.25 0.7 | 0 0 100 | 0 0 1 |
| | | 8 | 0 | 50 | 0 | 10 30 60 | 0.05 0.1 0.85 | 0 0 100 | 0 0 1 |
| | | 9 | 0 | 50 | 5 | 10 30 60 | 0.05 0.1 0.85 | 0 0 100 | 0 0 1 |
| | | 10 | 0 | 50 | 10 | 10 30 60 | 0.05 0.1 0.85 | 0 0 100 | 0 0 1 |
| | | 11 | 0 | 50 | 20 | 10 30 60 | 0.05 0.1 0.85 | 0 0 100 | 0 0 1 |
| | 70 | 1 | 21 | 60 | 0 | 0 0 100 | 0 0 1 | 0 0 100 | 0 0 1 |
| | | 2 | 21 | 40 | 0 | 0 0 100 | 0 0 1 | 25 45 30 | 0.15 0.3 0.5 |
| | | 3 | 21 | 5 | 0 | 0 0 100 | 0 0 1 | 10 40 50 | 0.05 0.25 0.7 |
| | | 4 | 21 | 5 | 0 | 0 0 100 | 0 0 1 | 10 20 70 | 0.05 0.1 0.85 |
| | | 5 | 10 | 70 | 0 | 0 0 100 | 0 0 1 | 0 0 100 | 0 0 1 |
| | | 6 | 10 | 70 | 0 | 25 45 30 | 0.15 0.3 0.5 | 0 0 100 | 0 0 1 |
| | | 7 | 7 | 70 | 0 | 10 40 50 | 0.05 0.25 0.7 | 0 0 100 | 0 0 1 |
| | | 8 | 0 | 70 | 0 | 10 20 70 | 0.05 0.3 0.65 | 0 0 100 | 0 0 1 |
| | | 9 | 0 | 70 | 5 | 10 20 70 | 0.05 0.3 0.65 | 0 0 100 | 0 0 1 |
| | | 10 | 0 | 70 | 10 | 10 30 60 | 0.05 0.3 0.65 | 0 0 100 | 0 0 1 |
| | | 11 | 0 | 70 | 20 | 10 30 60 | 0.05 0.3 0.65 | 0 0 100 | 0 0 1 |

**Supplementary Table 4: Stand sizes for random trees for biological datasets.** Three datasets D1, D2, D5 (in bold) does not have 100M trees stands. For the remaining nine datasets the majority of random trees (≥95) corresponding stands have a lower bound of 100M trees on stand size. In grey are datasets, for which all 100 random trees have a lower bound of 100M trees.

| ID | Internal ID | Species Group | Number of random trees with corresponding stand size | Stand size |
|---|---|---|---|---|
| **D1** | **d62_1027** | **Snails** | **100** | **1** |
| **D2** | **d103_852** | **Land plants** | **100** | **1** |
| D3 | d180_15 | Drosophilinae | 97 | 100M |
| | | | 1 | 24,310,125 |
| | | | 1 | 467,775 |
| | | | 1 | 118,125 |
| D4 | d237_74 | Carnivora | 100 | 100M |
| **D5** | **d267_20** | **Frogs** | **1** | **7** |
| | | | **5** | **5** |
| | | | **18** | **3** |
| | | | **76** | **1** |
| D7 | d279_27 | Primates-1 | 100 | 100M |
| D7 | d298_3 | Grasses | 100 | 100M |
| D8 | d372_79 | Primates-2 | 100 | 100M |
| D9 | d381_13 | Salamander | 98 | 100M |
| | | | 1 | 89,335,499 |
| D10 | d404_11 | Monocots | 100 | 100M |
| D11 | d435_18 | Sedges | 100 | 100M |
| D12 | d767_5 | Snakes | 95 | 100M |
| | | | 1 | 88,232,940 |
| | | | 1 | 64,480,248 |
| | | | 1 | 59,333,184 |
| | | | 1 | 28,704,375 |
| | | | 1 | 10,333,575 |
