## Supplementary figures and images for "Gentrius: identifying equally scoring trees in phylogenomics with incomplete data"

### Suppl.Fig.6-Strict_consensus_tree-D12-Snakes.pdf

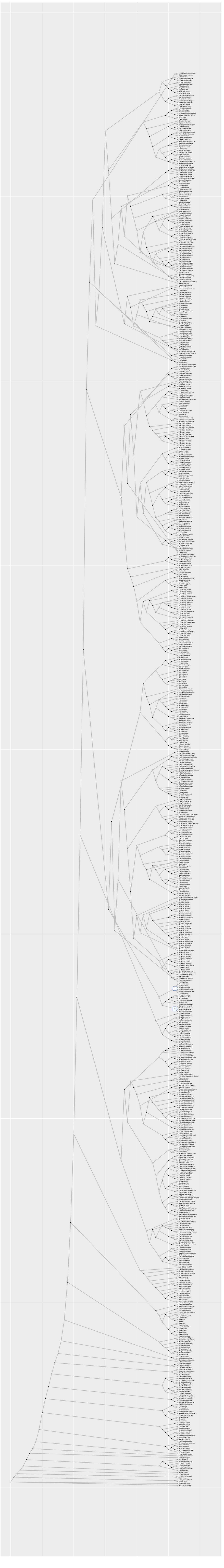

S: 315 | R: 99% | iB: 758 | mN: 2

### Suppl.Fig.7-Strict_consensus_tree-D6-Primates-1.pdf

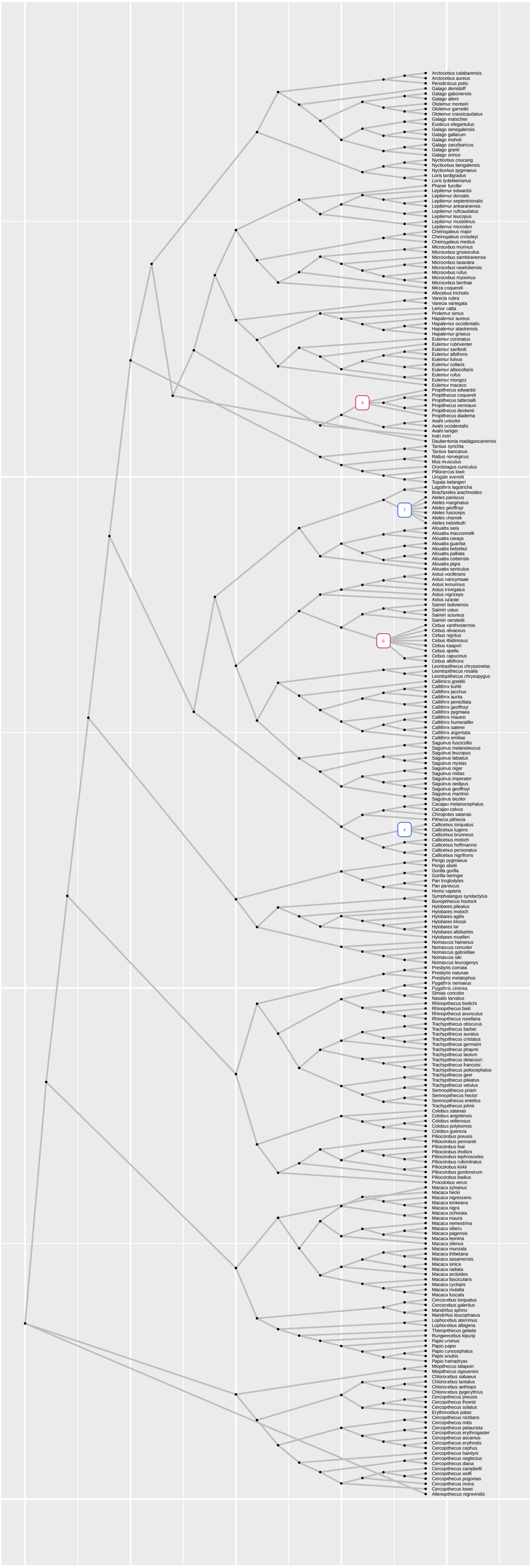


S: 9963 | R: 96% | iB: 265 | mN: 4

### Suppl.Fig.8-Strict_consensus_tree-D3-Drosophilinae.pdf

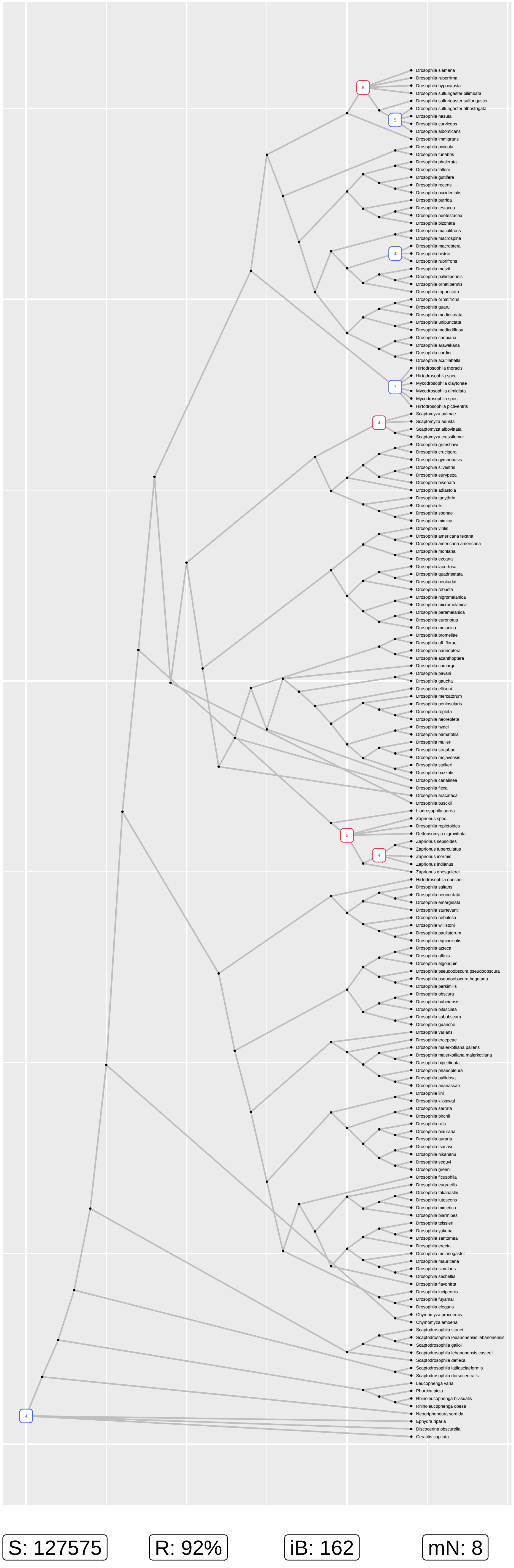

### Suppl.Fig.9-Strict_consensus_tree-D8-Primates-2.pdf

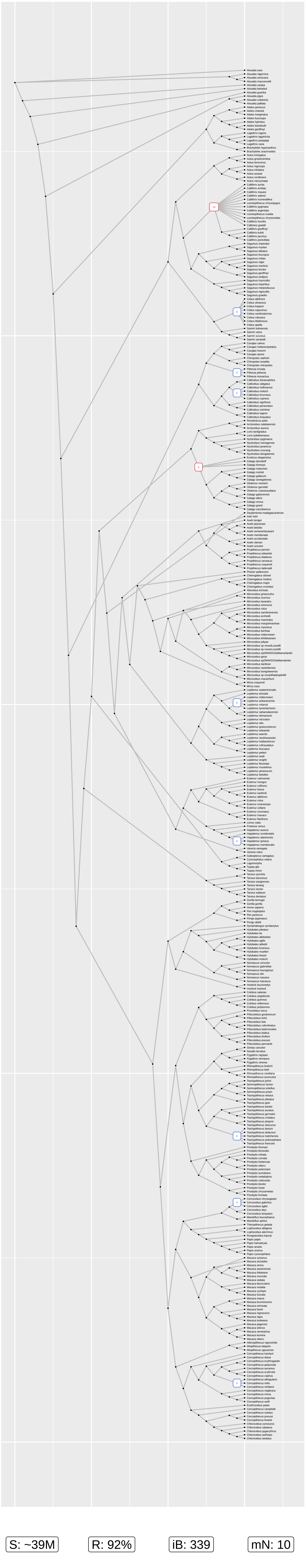

### Suppl.Fig.10-Strict_consensus_tree-D10-Monocots.pdf

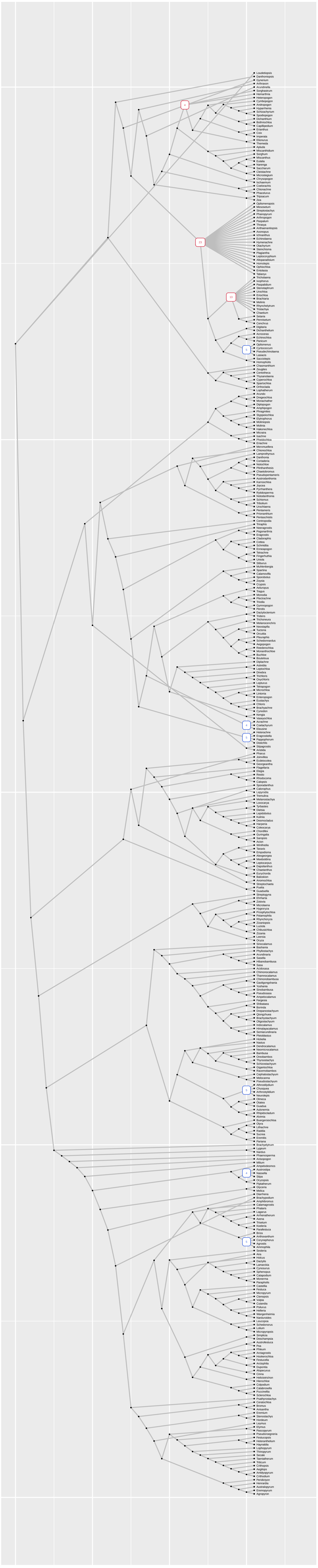

S: ~15M | R: 90% | iB: 360 | mN: 9

### Suppl.Fig.11-Strict_consensus_tree-D4-Carnivora.pdf

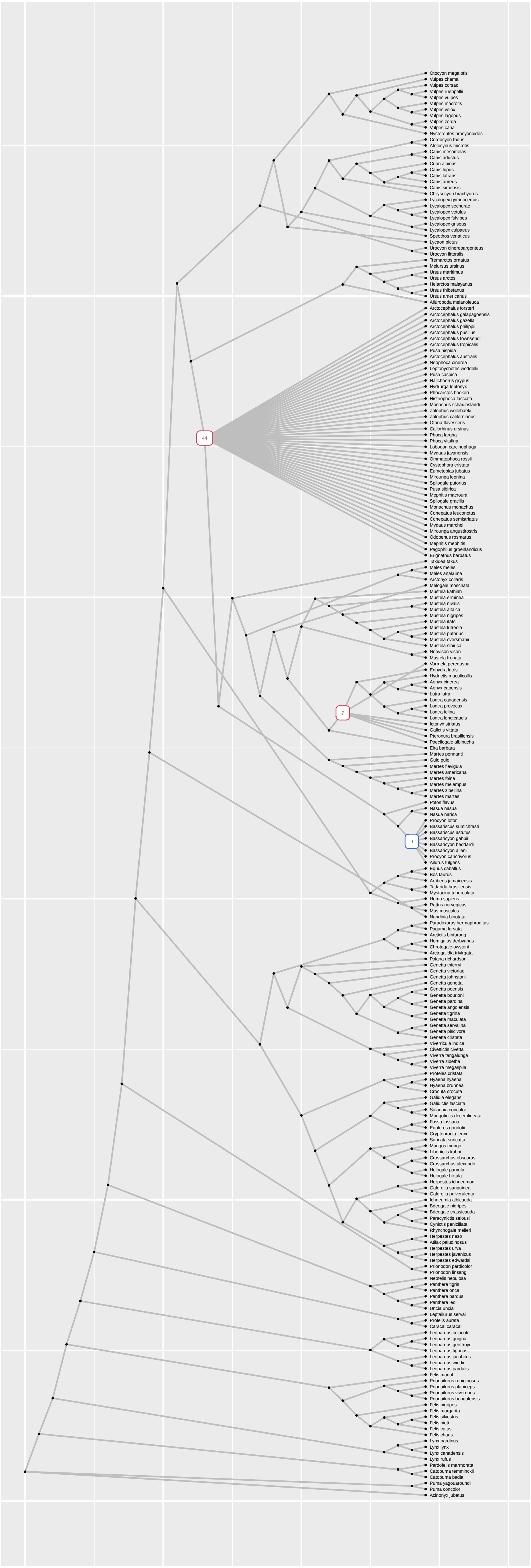


S: 29133

R: 78%

iB: 183

mN: 3
